# Neuronal activation affects the organization and protein composition of the nuclear speckles

**DOI:** 10.1101/2024.03.25.586583

**Authors:** Andrzej Antoni Szczepankiewicz, Kamil Parobczak, Monika Zaręba-Kozioł, Błażej Ruszczycki, Monika Bijata, Paweł Trzaskoma, Grzegorz Hajnowski, Dagmara Holm-Kaczmarek, Jakub Włodarczyk, Grzegorz Marek Wilczyński, Maria Jolanta Rędowicz, Adriana Magalska

## Abstract

Nuclear speckles, also known as interchromatin granule clusters (IGCs), are subnuclear domains highly enriched in proteins involved in transcription and mRNA metabolism and, until recently, have been regarded primarily as their storage and modification hubs. However, several recent studies on non-neuronal cell types indicate that nuclear speckles may directly contribute to gene expression as some of the active genes have been shown to associate with these structures.

Neuronal activity is one of the key transcriptional regulators and may lead to the rearrangement of some nuclear bodies. Notably, the impact of neuronal activation on IGC/nuclear speckles organization and function remains unexplored. To address this research gap, we examined whether and how neuronal stimulation affects the organization of these bodies in granular neurons from the rat hippocampal formation. Our findings demonstrate that neuronal stimulation induces morphological and proteomic remodelling of the nuclear speckles under both *in vitro* and *in vivo* conditions. Importantly, these changes are not associated with cellular stress or cell death but are dependent on transcription and splicing.

## 1. Introduction

The cell nucleus, although deprived of any internal membrane compartments, is far from being a homogeneous organelle. The structural elements of the nucleus are chromatin, organized into chromosomal territories and several types of nuclear bodies located within the interchromatin space. One kind of subnuclear body, which is very intensively studied are nuclear speckles. They were originally described with the use of light microscopy, first by Santiago Ramón y Cajal (Cajal, 1910) and later independently by J. Swenson Beck (Beck, 1961). Also, Hewson Swift with the use of electron microscopy demonstrated the presence of these structures within the nuclei (Swift, 1959). It is noteworthy that currently, two distinct terms describe these nuclear bodies, depending on the technique used for their original characterization. The term “nuclear speckles” is associated with light microscopy observations, and the name “interchromatin granule clusters” (IGCs) is related to electron microscopy visualization.

Nuclear speckles were initially described as a place of storage and modification of the factors involved in gene transcription, splicing, and mRNA export (Spector and Lamond, 2011). However, several reports indicate that nuclear speckles could also play a more dynamic role in these processes. Numerous groups have shown that nuclear speckles are associated with loci of several active genes (Brown et al., 2008; Brown et al., 2006; Jolly et al., 1999; Moen Jr et al., 2004; Nielsen et al., 2002; Smith et al., 1999; Szczerbal and Bridger, 2010; Xing et al., 1995). Furthermore, the genome-wide studies using SPRITE (Split-Pool Recognition of Interactions by Tag Extension) and TSA-Seq confirmed that certain chromosomal regions exhibit preferential and deterministic localization to the proximity of the nuclear speckles (Chen et al., 2018; Quinodoz et al., 2018), and it appears to be conserved among different cell types (Zhang et al., 2021). It has been also shown that genes localized in the vicinity of nuclear speckles demonstrate higher spliceosome concentrations, increased spliceosome binding to their transcripts, and higher co-transcriptional splicing levels than genes located further away from nuclear speckles (Bhat et al., 2023). Moreover, the presence of several gene transcripts has been shown within or at the edges of the nuclear speckles (Hattinger et al., 2002; Johnson, 2000; Melcak et al., 2000; Shopland et al., 2002; Smith et al., 1999). Also, it was shown that microinjected pre-mRNAs containing functional splicing sites were stored and spliced within nuclear speckles (Dias et al., 2010). In line with this observation, Carvalho et al. showed that splicing arrest resulted in the accumulation of the intron-containing transcripts in the nuclear speckles (Carvalho et al., 2017). Another observation using APEX2 labelling showed that nuclear speckles are highly enriched in transcripts harboring retained introns (Barutcu et al., 2022). Alongside these observations, a phosphorylated form of SF3b155 protein, specific to the catalytically active spliceosomes, was detected in nuclear speckles (Girard et al., 2012). Overall, the above insights provide strong evidence supporting the claim, that some of the splicing events take place in and/or at the edge of the nuclear speckles. In addition, the role of nuclear speckles was linked to mRNA export: association of mRNA transcripts of intron-less genes with nuclear speckles enhances their export by facilitating the recruitment of the multiprotein TREX (TRanscription-EXport) complex (Wang et al., 2018). Considering the above, the concept of nuclear speckles as versatile hubs, connecting mRNA transcription, splicing, and export (‘gene expression hubs’ as proposed by Hall et al.) appears to be correct and appealing (Hall et al., 2006).

The above-mentioned studies were carried out on transformed and/or proliferating cells. However, very little is known about the function of IGC/nuclear speckles in the nuclei of terminally differentiated cells such as neurons. In neurons of rat trigeminal ganglion, a correlation between the cell and nuclear speckles size has been observed. Larger neurons with high transcriptional activity have smaller nuclear speckles and show a diffusive snRNP (small nuclear ribonucleoproteins) pattern in the nucleoplasm, while smaller neurons, with low transcriptional activity, have larger nuclear speckles (Pena et al., 2001). In line with these results, a reversible change in the nuclear speckles organization in response to the damage to the axonal termini was reported. The enlarged nuclear speckles are more abundant in U2 snRNP B (the component of spliceosome) and the authors connect this observation with the lower transcription activity. The axonal damage impaired the recruitment of splicing factors to the pre-mRNA splicing sites and increased the concentration of factors stored in the nuclear speckles (Navascues et al., 2004).

Our preliminary data showed that *in vitro* neuronal stimulation increased the number of nuclear speckles (Grabowska et al., 2022). To further investigate the function of nuclear speckles in the rat hippocampal neurons from the dentate gyrus, we decided to expand these studies by examining the effects of neuronal stimulation by the *in vivo*-induced seizures on the speckle morphology and protein composition.

## 2. Materials and methods

### 2.1. Animals

The experiments were carried out on the *Rattus norvegicus* Wistar strain. Adult males, 8-9 weeks old, and 200-250 g were used in the seizure experiments. P7 rat neonates were used for hippocampal organotypic culture preparation. In both cases, the animals came from the Mossakowski Institute of Experimental and Clinical Medicine PAS in Warsaw. The adult animals were housed in group cages under a 12-hour light-dark cycle with unlimited access to water and food. All animal procedures and experimental conditions were approved by the Local Ethics Committee for Animal Experiments (consent no. 774/2015).

### 2.2. Induction *status epilepticus* with kainic acid

*Status epilepticus* was induced by 2-4 intraperitoneal injections of 5 mg/kg kainic acid (Milestone Pharmtech USA Inc., 6M-0100) in 1 h intervals (Hellier et al., 1998). The control group was administered with 4 doses of the saline solution according to the same schedule. After each injection, rats were placed in separate cages and the development of the seizures was observed. Seizure assessment was made based on the modified Racine Scale (Ben-Ari, 1985; Racine, 1972) as described in (Skupien-Jaroszek et al., 2021). When grade 5 or higher seizures occurred at least 3 times per hour, kainate injections to the animal were terminated; most of such animals developed full-*status epilepticus*. Rats were sacrificed 2 h from the beginning of *status epilepticus* as well as 2 h and 24 h, and 1 week after injection termination. Rats allocated for the microscopic experiments were anesthetized by intraperitoneal administration of Morbital (pentobarbital sodium and pentobarbital, Biowet Puławy sp. z o.o) at a dose of 1.5 ml/kg and subjected to transcardial perfusion, first with a PBS containing 10 U/l heparin, and then with a fixative appropriate for the further procedures. Animals allocated for the mass spectrometry were anesthetized by Aerrane inhalation (isoflurane, Baxter International Inc.). Control rats were anesthetized 4.5 h after the saline injection, which corresponds to the mean time between the first dose of kainic acid and the sacrification of the stimulated animals.

### 2.3. Induction seizures with pentylenetetrazole

Pentylenetetrazole was administered intraperitoneally in a single dose of 50 mg/kg. Injected animals were kept in a styrofoam box to minimize injuries that may occur as a result of strong seizures (Obay et al., 2007). Seizures started usually a few minutes after the proconvulsant administration and lasted for several minutes. Two hours after the beginning of seizures, rats were sacrificed and perfused as described above.

### 2.4. Induction of chemically evoked LTP (cLTP), transcription, and splicing inhibition in rat hippocampal organotypic slice cultures

Organotypic slice culture was prepared as previously described (Stoppini et al., 1991). Briefly, the hippocampus was isolated from 6-day-old rats in ice-cold Gey’s Balanced Salt Solution with 28 mM glucose. Using a McIlwain tissue chopper, 300 µm-thick sections, perpendicular to the long axis of the hippocampus, were prepared. 3-4 sections were placed on one Millicell insert’s membrane. Inserts were then placed in 6-well plates with pre-incubated growth medium (Minimum Essential Medium with 25% Hank’s Balanced Salt Solution, 25% horse serum, 1% Penicillin-Streptomycin, 1 mM GlutaMax, and 28 mM glucose). Cultures were grown at 37°C under an atmosphere containing 5% CO_2_ and the medium was exchanged every 4 days.

After approximately 2 weeks of culture, neurons were subjected to the chemically induced LTP (cLTP) (Szepesi et al., 2014) by the addition of 50 µM forskolin, 0.1 µM rolipram, and 50 µM picrotoxin to the medium. After 2-h stimulation, cultures were fixed. Transcription inhibition was achieved by the addition of 1 h before the stimulation of 1 µg/ml actinomycin D to the medium. To block pre-mRNA splicing 10 µM pladienolide B was used for 2 and 8 h prior to the cLTP. Control samples were treated with the appropriate volume of the dissolvent (DMSO, concentration not higher than 1%).

### 2.5. Immunofluorescence labelling and confocal microscopy imaging

Perfusion was performed with a 4% solution of paraformaldehyde in PBS at room temperature. The extracted brains were post-fixed for 12 h in the same solution at 4°C, then rinsed briefly with PBS, and cryoprotected by saturation with 30% sucrose in PBS. Such brains were then frozen in heptane at −80°C and stored at this temperature until dissection. Next, they were cut into 40 µm thick coronal sections with cryostat and stored for further use at −20°C in an anti-freezing medium (30% glycerol, 30% ethylene glycol, 10 mM phosphate buffer pH 7.4). Immunofluorescence labelling was performed in the free-floating sections containing dorsal hippocampal formation (Bregma −3.30 mm to −4.30 mm). Before the labelling, sections were permeabilized for 10 min in 0.5% Triton X100 in PBS and subjected to nonspecific protein binding quiescence for 60 min in 5% normal donkey serum (NDS) in PBS with 0.1% Triton X100 (PBS-T). Such slices were incubated with mouse anti-SC35 primary monoclonal antibody (1:200, Abcam PLC, ab11826) overnight at 4°C. After washing with PBS-T, Cy3 AffiniPure F(ab’)₂ Fragment Donkey Anti-Mouse secondary antibody (1:200, Jackson ImmunoResearch Ltd, 715-166-150) was applied in 5% NDS in PBS-T for 2 h at room temperature. Subsequently, cFos protein was labelled in the same manner using rabbit polyclonal antibody c-Fos (9F6) (1: 1000, Cell Signaling Technology Inc, 2250) and Alexa Fluor 488 AffiniPure F(ab’)₂ Fragment Donkey Anti-Rabbit (1:200, Jackson ImmunoResearch Ltd, 711-546-152) as a secondary antibody. Cell nuclei were labelled with Hoechst 33342 or TO-PRO-3 in PBS, for 10 min. After a triple 10 min wash in PBS, sections were mounted on slides in Vectashield medium.

Confocal images was taken with a Leica TCS SP8 confocal microscope (Leica Microsystems GmbH, Germany), equipped with a WLL 470-670 nm and 405 nm diode laser, HC PL APO CS2 63x/1.40 Oil lens, with a pixel size of ∼ 70 nm in the XY, and ∼210 nm in the Z plane, with sequential (each channel separately) double averaging of the scanned lines. Imaging was performed in the Laboratory of Imaging Tissue Structure and Function, Nencki Institute of Experimental Biology, PAS.

### 2.6. Nuclear speckles 3D morphology analysis

The 3D morphological analysis of nuclear speckles was performed using custom-written software utilizing a continuous boundary tracking algorithm, which enables the separation of the closely packed nuclei of granule cells in the dentate gyrus (DG) (Ruszczycki et al., 2019). Each nucleus was saved as a separate TIF file. Next using a 3D Object Counter ImageJ Fiji plug-in, the number, volume, and area of nuclear speckles for individual nuclei were determined. Based on the obtained values of the specklès area (S) and volume (V), its form factor (ff) was determined using the following formula:

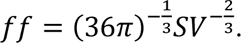

The form factor coefficient for a sphere has the value of 1 and increases as the object becomes less spherical.

### 2.7. Transmission electron microscopy

Animals were perfused with a fixative solution containing 4% paraformaldehyde and 2.5% glutaraldehyde in PBS. Extracted brains were additionally fixed for 12 h in the same solution at 4°C. After washing in PBS, brains were dissected into 100 µm thick coronal sections with VT 1000 S vibratome. Sections containing dorsal hippocampus (Bregma −3.30 mm to −4.30 mm) were stored in PBS with 0,1% sodium azide at 4°C for further use.

Hippocampal organotypic slice cultures were fixed with the same fixative, added first onto the membranes of the Millicell inserts and then also to the culture dish well in a place of the removed medium. After 1-h fixation, slices were rinsed 3 x 10 min in PBS and stored in the same medium with 0,1% sodium azide at 4°C. Prior to further preparation, slices were gently peeled off the membrane with a brush.

The next steps in the procedure were the same for both the brain sections and the organotypic culture slices. Nuclear speckles were contrasted by incubation with bismuth salts according to Lock and Huie’s method (Locke and Huie, 1977). Briefly, brain sections or organotypic slices were washed 3 times, 10 min each in 0.1 M triethanolamine, then in a bismuth salt solution prepared according to the original recipe (solution A containing 400 mg sodium tartrate in 10 ml of 1 N NaOH added dropwise to 200 mg bismuth(III) nitrate and solution B containing 0.2 M triethanoloamine mixed in the ratio 1:3, pH 7.0) for 1.5 h and washed as previously, first with 0.1 M triethanolamine and then with a phosphate buffer. After the final wash, the tissue was incubated in a 1% osmium tetroxide solution and washed 3 x 10 min in water. Dehydration was carried out using water solutions with an increasing ethanol concentration: 50%, 70%, 90%, 96% (10 min each), and finally twice with pure ethanol (20 min each). Dehydrated tissue was incubated first with a 1:1 ethanol and propylene oxide solution (10 min), which was then exchanged for pure propylene oxide (twice, 15 min each). Embedding with Epoxy resin was initiated with a mixture of propylene oxide and resin in a 1:1 ratio, exchanged for 1:2 solution (1 h each), and finally incubated twice in pure resin, first for 2 h and then overnight. On the next day, the tissue in a new portion of resin was placed between two sheets of Aclar foil and polymerized at 60°C for two days. After removing the Aclar foil, a fragment of the dentate gyrus was cut out and glued with cyanoacrylate glue to a blank resin block. The resulting 60 nm ultra-thin sections were cut with Leica Ultracut R ultramicrotome and collected on TEM grids. Uranyl acetate and lead citrate staining were omitted. Images of the DG granular cells nuclei were obtained using a JEM 1400 transmission electron microscope (JEOL Ltd., Japan) equipped with an 11-megapixel Morada G2 camera (EMSIS GmbH, Germany) in the Laboratory of Electron Microscopy of the Nencki Institute of Experimental Biology PAS.

A relative volume of IGCs aggregates was determined on electron micrograph according to principle which states that relative area occupied by the cross-section of a smaller object inside the cross-section of a larger object is a determinant of the relative volume occupied by the smaller object in the larger object (Elias and Hyde, 1980).

### 2.8. Serial Block-Face Scanning Electron Microscopy and 3D IGC/nuclear speckle reconstruction

A fragment of DG prepared according to the above procedure was glued to a metal pin with CircuitWorks Conductive Epoxy glue (Chemtronics, CW2400). After trimming, the edges of the block were coated with a conductive paste (Ted Pella Inc., 16062). Sigma VP 3View scanning electron microscope (Carl Zeiss AG, Germany) equipped with a Gatan 3View module (Gatan Inc., USA) and diamond knife was used to collect a stack of images with resolution 5.8 nm x 5.8 nm in YX, and 50 nm in Z plane (Z = thickness of cutting sections). 3D reconstruction was performed with the Imaris 7.4.2 software (Bitplane AG, Switzerland). The IGC/nuclear speckles were reconstructed using automatic boundary determination by absolute intensity. The surface of the cell nuclei was reconstructed by manually drawing boundaries at the site of the nuclear envelope.

### 2.9. High-pressure freezing and freeze substitution

The perfusion of animals was performed with a fixative solution containing 4% paraformaldehyde and 0.1% glutaraldehyde in 0.1 M phosphate buffer. Brains were cut and stored in the same way as for the TEM procedure. Brain sections were cryoprotected by 20 min incubation in 30% sucrose in PBS at 4°C. A 1.5 mm in diameter piece of the dentate gyrus was cut out from the slices with a specimen punch and placed on membrane carriers (Leica Microsystems GmbH, 707898) coated with lecithin (1% solution in chloroform). The space between the tissue and the membrane carriers was filled with hexadecane. The rapid loader was used to load the samples into the EM PACT2 (Leica Microsystems GmbH, Austria), where they were frozen at −196°C and 2000 bar. After that, samples were transferred in a liquid nitrogen bath to an EM AFS2 machine (Leica Microsystems GmbH, Austria), equipped with an automatic EM FSP processor. Briefly, a frozen piece of tissue, resting on the membrane carrier, was incubated in a 1.5% uranyl acetate solution in methanol cooled to −90°C for 41 h. Then it was rinsed 3 times for 10 min in pure methanol and incubated in the mixture of methanol and Lowicryl (1:1 and 1:2, 2 h each, all steps at −45°C). Next, pure Lowicryl was provided for 12 hours. After replacing the resin with a fresh one, UV polymerization was carried out for 24 h at −45°C. UV was continued for 9 h while the temperature was rising to 0°C and then for the next 35 h. Upon completion, the resin blocks were removed from the container, the membrane carriers separated and the block of resin was trimmed. 60 nm-thick ultrathin sections were obtained on a Leica Ultracut R ultramicrotome. Sections were placed on nickel grids coated with formvar and carbon. A detailed freeze substitution procedure on an EM AFS2 machine equipped with an automatic EM FSP processor is presented in Tab. S1.

**Table S1.** Freeze substitution procedure on EM AFS2 machine equipped with an automatic EM FSP processor.

| Step | Initial temp. | Final temp. | Duration h:min | Reagent | Action of EM FSP | UV | Comments |
| --- | --- | --- | --- | --- | --- | --- | --- |
| 1 | - 90°C | - 90°C | 30:00 | 2% uranyl acetate in methanol | none | no | dehydration and contrasting |
| 2 | - 90°C | - 45°C | 11:15 | 2% uranyl acetate in methanol | none | no | dehydration and contrasting |
| 3 | - 45°C | - 45°C | 00:10 | 100% methanol | removes and fills | no | dehydration and washing |
| 4 | - 45°C | - 45°C | 00:10 | 100% methanol | removes and fills | no | dehydration and washing |
| 5 | - 45°C | - 45°C | 00:10 | 100% methanol | removes and fills | no | dehydration and washing |
| 6 | - 45°C | - 45°C | 02:00 | 100% methanol: Lowicryl 1:1 | removes and fills | no | embedding |
| 7 | - 45°C | - 45°C | 02:00 | 100% methanol: Lowicryl 1:1 | removes and fills | no | embedding |
| 8 | - 45°C | - 45°C | 02:00 | Lowicryl | removes and fills | no | embedding |
| 9 | - 45°C | - 45°C | 10:00 | Lowicryl | removes and fills | no | embedding |
| 10 | - 45°C | - 45°C | 00:01 | Lowicryl | removes and fills | no | fresh portion of resin for polymerization |
| 11 | - 45°C | - 45°C | 24:00 | Lowicryl | none | yes | polymerization |
| 12 | - 45°C | 0°C | 09:00 | Lowicryl | none | yes | polymerization |
| 13 | 0°C | 0°C | 35:00 | Lowicryl | none | yes | polymerization |

### 2.10. Immunogold protein labelling

The immunogold labelling procedure was performed with the “on a drop” method, using the modified protocol provided by Mathiisen et al. (Mathiisen et al., 2006). The protocol includes: (1) 10 min blocking of the free aldehyde groups in a 50 mM glycine solution in TBST buffer (Tris-buffered sodium chloride with 0.1% Triton X-100), (2) blocking non-specific protein interactions with 2% BSA in TBST for 10 min, (3) incubation with the primary antibody in 2% BSA in TBST for 2 h at room temperature or overnight at 4°C, (4) two 10 min washes in TBST proceeded by triple few seconds rinse. The solution remaining on the sample was removed from the grid with filter paper between the washes. Subsequently, the sample was (5) incubated for 2h in a 1:10 solution of colloidal gold-conjugated secondary antibody in TBST with 2% BSA and 0.05% polyethylene glycol PEG-2000, (6) rinsed 6 times with deionized water (remaining water was, as previously, removed from the grid) and (7) dried. Before the labelling, the specificity of the above procedure was determined by utilizing non-specific rabbit and mouse primary antibodies that showed no labelling on the tested material.

A list of the primary antibodies used includes: mouse monoclonal IgM anti-phospho-SR (3C5) (1:20, gift from David Spector, Cold Spring Harbor Laboratory, NY, USA); rabbit polyclonal IgG anti-RNA polymerase II CTD repeat YSPTSPS phospho S5 (1:20, Abcam PLC, ab5131); rabbit polyclonal IgG anti-RNA polymerase II CTD repeat YSPTSPS phospho S2, (1:20, Abcam PLC, ab5095), rabbit polyclonal IgG anti-coilin (H-300) (1:20, Santa Cruz Biotechnology Inc., sc-32860), rabbit anti-PSF (H-80) polyclonal IgG (1:20, Santa Cruz Biotechnology Inc., sc-28730), rabbit anti-Sam68 polyclonal IgG (C-20) (1:20, Santa Cruz Bio - technology Inc., sc-333), mouse anti-SF3a66 monoclonal IgG (4G8) (1:20, Abcam PLC, ab77800). List of the secondary gold-conjugated antibodies used: donkey anti-rabbit IgG (H&L) 10nm (1:10, Aurion, 110.311), donkey anti-mouse IgG (H&L) 10nm (1:10, Aurion, 110.322).

The labelling density (number of gold particles/µm^2^) was determined by counting the gold particles within the IGCs profiles and relating this value to the area of this profile (Griffiths, 1993). The surface area of the profiles and the number of gold particles were measured in Fiji ImageJ (Schindelin et al., 2012) using the Cell Counter plugin.

### 2.11. Localization of DNA at the ultrastructural level

DNA immunolocalization at the ultrastructural level was performed according to Marc Thiry’s method (Thiry, 1992), in which the terminal deoxynucleotide transferase (TdT) adds labelled nucleotides (dUTP) to the DNA fragments protruding from the resin surface of an ultrathin section. The labelling was performed on ultrathin sections prepared as described in the chapter ‘High-pressure freezing, freeze substitution, and immunogold’. The protocol included: (1) 1h incubation at 37°C with a reaction mixture containing 600 U/ml TdT polymerase (Thermo Fisher Scientific Inc., 10533-065) and 7.5 µM fluorescein-labelled dUTP (Thermo Fisher Scientific Inc., R0101), (2) rinsing in deionized water 6 times, (3) 10 min blocking sites of non-specific protein binding with 2% BSA in TBST, (4) 2h incubation with mouse anti-fluorescein antibody conjugated with 10 nm colloidal gold (1:10, Aurion, 810.233), (5) 6 rinses in deionized water and (6) drying. Samples were imaged with a JEM 1400 TEM as described before. As a control, incubation with anti-fluorescein antibody conjugated with colloidal gold was performed, omitting the TdT reaction step. No non-specific binding of the antibody under these conditions was detected.

### 2.12. Detection of neurodegeneration in the hippocampus

Neurodegeneration in hippocampal cells was detected with Fluoro-Jade B staining (Millipore, AG310), according to the manufacturer’s protocol. Staining was performed on tissue sections taken one day and one week after the end of the *status epilepticus*. Imaging was carried out using a Leica DMI6000 microscope with an I3 filter allowing the excitation wavelength of 450-490 nm at which Fluoro-Jade B emits green light with an emission peak at 525 nm.

### 2.13. Detection of apoptosis in the hippocampus by TUNEL method

Click-iT TUNEL Alexa Fluor 488 Imaging Assay (Thermo Fisher Scientific Inc., C10245) was used to detect apoptosis in the hippocampal neurons as a result of the *status epilepticus*. The manufacturer’s original protocol is designed to detect apoptosis in adherent cells, however, we successfully adapted it for brain cryostat sections. Sections were washed in PBS from an anti-freezing medium. The next steps were made according to the manufacturer’s procedure except that reaction and labelling were performed on the hippocampal part of the brain section and the final volume of the reaction was adjusted to 100 µl. The imaging was carried out on a Leica DMI6000 microscope with an I3 filter enabling the excitation wavelength of 450-490 nm, at which Alexa Fluor 488 emits green light with an emission peak at 519 nm.

### 2.14. Proteomic analysis of nuclear fraction enriched in IGC/nuclear speckles by mass spectrometry

For technical reasons, the nuclear fraction enriched in IGC/nuclear speckles was obtained from the whole hippocampus.

#### 2.14.1. Enrichment of the nuclear fraction with IGC/nuclear speckles

Two hours after the end of *status epilepticus*, rats were sacrificed by inhalation of a lethal dose of isoflurane and decapitated. Brains were removed from the skull and meninges, and the hippocampi were isolated on ice. Next, the hippocampi were frozen in liquid nitrogen and stored at −80°C until further use. Hippocampi were obtained from 60 rats with *status epilepticus* and 60 control animals.

Nuclei isolation was performed according to Matevossian and Akbarian’s method (Matevossian and Akbarian, 2008) with minor modifications. Each hippocampus was homogenized on ice with a Dounce homogenizer in 5 ml of tissue lysis buffer (0.32 M sucrose, 5 mM CaCl_2_, 3 mM MgCl_2_, 0.1 mM EDTA, 0.1% Triton X-100, 1x Sigmafast Protease Inhibitor, 10 mM Tris-HCl). Next, the homogenate was purred over 6.3 ml sucrose solution (1,8 M sucrose, 5 mM CaCl_2_, 1x Sigmafast Protease Inhibitor, 10 mM Tris-HCl) in 13.2 ml centrifuge tubes (Beckman Coulter Inc., 344059) and centrifuged for 2.5 h at 30000 RPM (∼154 000 g, 4°C). The supernatant was removed and the nuclei pellet was resuspended in TM5 buffer (10 mM Tris-HCl, 5 mM MgCl_2_, pH 7.4) with Sigmafast Protease Inhibitor. About 1 300 000 nuclei per hippocampus were isolated. The nuclei were resuspended in the nuclear lysis buffer (TM5 buffer containing 1% Triton X-100, 2 mM Ribonucleoside Vanadyl Complex, and a mixture of protease inhibitors); the sample was incubated for 5 min on ice and then frozen in liquid nitrogen. Such samples were stored at −80°C until the further steps.

The next steps were performed according to Mintz et al. (Mintz et al., 1999). After thawing, nuclear lysates from 40 hippocampi (20 rats) were pooled, centrifuged, and resuspended in TM5 buffer with protease inhibitors. RNAse-free DNase I was added to a concentration of 10 U/µl and samples were incubated at 4°C for 1 h with slow mixing. The digested chromatin was removed by adding 5 M NaCl to a final concentration of 0.5 M. The samples were incubated for 5 min on ice, centrifuged (780 g, 5 min, 4°C), and the supernatant was discarded. This step was repeated three times. The final pellet was resuspended in a 100 µl mixture of 95.5 µl of 0.5 M NaCl solution and 0.5 µl of 1 M DTT and left for 5 min on ice. The suspension was passed 10 times through a 0.33 mm (29 G) needle, then homogenized in a 1 ml Dounce glass homogenizer on ice, using a tight piston. The homogenate was transferred to a tube containing 500 µl of 0.25 M Cs_2_SO_4_ and centrifuged for 2 min at 20800 g. The supernatant containing the nuclear fraction enriched in IGC/nuclear speckles was transferred to a new tube to which 200 µl of TM5 buffer was added. The suspension was placed in 0.5 ml centrifuge thick-walled tubes (Thermo Fisher Scientific Inc., 45235-AV) and centrifuged for 1 h at 157 000 g and 4°C. The supernatant was discarded and the pellet, containing the nuclear fraction enriched in IGC/nuclear speckles, was resuspended in 60 µl of TM5 buffer and frozen at −80°C until further analysis.

#### 2.14.2. Validation of nuclear fraction enrichment in IGC/nuclear speckles

Speckle pellets were sonicated on ice with two 10-second pulses of Branson S-250D sonicator set to 30% of the maximum capacity, intermitted by a 10-second cooling interval. Protein concentration was measured using the Pierce BCA Protein Assay reagent kit (Thermo Fisher Scientific Inc., 23227). Protein electrophoretic separation was performed using the Laemmli method (Laemmli, 1970). After the electrotransfer of proteins onto the nitrocellulose membrane, chemiluminescence immunodetection was used to detect proteins. A rabbit anti-SFRS2 (SC35) antibody (1: 500, Proteintech Group Inc., 20371-1-AP) and Peroxidase AffiniPure Mouse Anti-Rabbit IgG (H + L) antibody (1: 5000, Jackson ImmunoResearch Ltd., 211-035-109) were used. Visualization of the antibody binding sites was achieved using Amersham ECL Prime Western Blotting (GE Healthcare UK Ltd, RPN2232) according to the manufacturer’s protocol.

#### 2.14.3. Mass spectrometry

The fraction enriched in IGC/nuclear speckle proteins was dissolved in 50 mM Tris-HCl buffer with 150 mM NaCl, pH 8.5, and reduced for 1h with 5 mM TCEP at 60°C. Thiol groups were blocked for 45 with 10 mM iodoacetamide min at 25°C. The proteins were then digested overnight with 10 ng/μl trypsin solution. Finally, samples were subjected to spectrometric analysis at the core facility of the Institute of Biochemistry and Biophysics PAS, Warsaw.

The MS/MS data were analysed using the Mascot Distiller software. The obtained results were compared with the SwissProt protein database (Swissprot 2018_02;17,205 sequences) using the Mascot Server 2.4.3 algorithm. The following application search parameters were used: enzyme - trypsin; the number of sites omitted during trypsin cleavage - 1; permanent protein modifications - carabmidomethylation (C); variable modifications - oxidation (M); tool - HCD; options Decoy - active, animal - *Rattus norvegicus*. The FDR (false discovery rate) was estimated using Mascot Decoy and was kept below 1%. Then, merged lists of peptides identified in the LC-MS/MS runs in all replicates for a given experimental setup were created in the MScan program. The RAW data obtained from the LC-MS traces were converted to a format that can be used by the MSparky program using the MSconverter program. In the conversion process, the LC-MS data are transformed into two-dimensional matrices. One of the dimensions of the matrix corresponds to the signals of the daughter ions that appeared in the chromatographic run (mass-to-charge ratio m/z). The MSparky program allows the assignment of the signals on the 2D maps to the appropriate peptides from the list created by the MScan program. The assignment was performed using the “peak-picking” method using information about their m/z value, retention time, or average peptide mass. In the data analysis, the 0/1 criterion was adopted, where 0 means no peptide on the map, and 1 - its presence. When verifying the correctness of the peptide assignments, the following criteria were taken into account: m/z value deviation, retention time deviation, EMRSE (Envelope Root Mean Squared Error), i.e. the discrepancy between the expected isotope envelope of the peak and the experimental envelope. Verification of the experimental data required manual checking of each peptide assignment to the isotopic envelope.

#### 2.14.4. The functional and quantitative analysis of proteins

The functional analysis of the identified proteins was performed based on the UniProt database. Since many of the identified proteins have more than one function, they were classified into several functional categories simultaneously. The sum of the number of proteins in the lower functional categories (e.g., splicing, mRNA export, etc.) is therefore greater than the total number of proteins in the superior category (e.g., mRNA metabolism). For the same reason, the sum of the number of proteins in all categories is greater than the total number of categorized proteins.

The quantitative analysis was performed using the Diffprot program (Malinowska et al., 2012; Zareba-Koziol et al., 2019). The ratio of protein concentrations in the compared groups, i.e. in the control group and the group with *status epilepticus* was determined. The fold change was presented as log_2_ of the ratio of the peptide concentrations in the control versus the *status epilepticus* animals. This parameter assumes negative values in case of a decrease and positive values in case of an increase in protein level. The statistical significance of protein concentration ratios was determined using a non-parametric permutation test, which determines the distribution of test statistics with the null hypothesis being true, based on random changes in the assignments of samples to groups and peptides to proteins. The resulting q values were adjusted for multiple testing to control FDR.

### 2.15. Statistical analysis of the results

The results are presented as arithmetic means with standard errors (presented in the text as mean value ± error value). The normality of the distributions was validated by the Kolmogorov-Smirnov test or - if the sample size allowed - by the Shapiro-Wilk test. The significance of differences between the groups was tested with the Student’s t-test or with the Mann-Whitney U test. The standard value of p <0.05 was adopted as the critical level of significance. The significance level was marked in the graphs as: * for p <0.05, ** - p <0.01, *** - p <0.001, and **** - p <0.0001. In the case of no significance, the abbreviation ns was used. The number of biological replicates (animals or organotypic cultures) was given in the figure description as N. The number of nuclei or nuclear cross-sections was given in the figure description as n.

## 3. Results

### 3.1. The morphology of the nuclear speckles changes upon neuronal activation

To test whether neuronal activation affects the organization of the IGC/nuclear speckle, we used one of the rat models of epileptic seizures. In this model, the intraperitoneal administration of kainic acid, an analog of glutamate, induces *status epilepticus* accompanied by strong activation of dentate gyrus (DG) granule neurons (Ben-Ari, 1985; Sperk, 1994) and increases expression of immediate early genes (Zagulska-Szymczak et al., 2001).

First, using confocal fluorescence microscopy combined with 3D quantitative measurements we examined morphological features of nuclear speckles in the DG granule neurons. Nuclear speckles were stained with an antibody recognizing SC35 (SRSF2), one of the major SR proteins present in these structures. The stimulation of granular neurons was confirmed by the detection of the c-Fos transcription factor, which expression accompanies neuronal activation (Dragunow and Robertson, 1987; Morgan et al., 1987) (Fig. 1A). Contrary to the *in vitro* model studied previously (Grabowska et al., 2022), we did not observe statistically significant changes in the mean number and mean volume of nuclear speckles (Fig. 1B and 1C). Instead, we noticed differences in the frequency distribution of the smallest (≤1.5 µm^3^), and the largest (≥9.5 µm^3^) nuclear speckles, which were more abundant upon stimulation (Fig. 1D). Additionally, very large nuclear speckles (in the range of 13.5 µm^3^ - 18.5 µm^3^), although not numerous, occurred only in the nuclei of stimulated cells (Fig. 1D). Moreover, upon activation, nuclear speckles with a volume ≥4.75 µm^3^, become more spherical than nuclear speckles of the same size in the control samples (Fig. 1E). In both cases (control and stimulation), the nuclear speckles showed the same typical localization in the interchromatin space (Fig. 1A).

**Figure 1.**
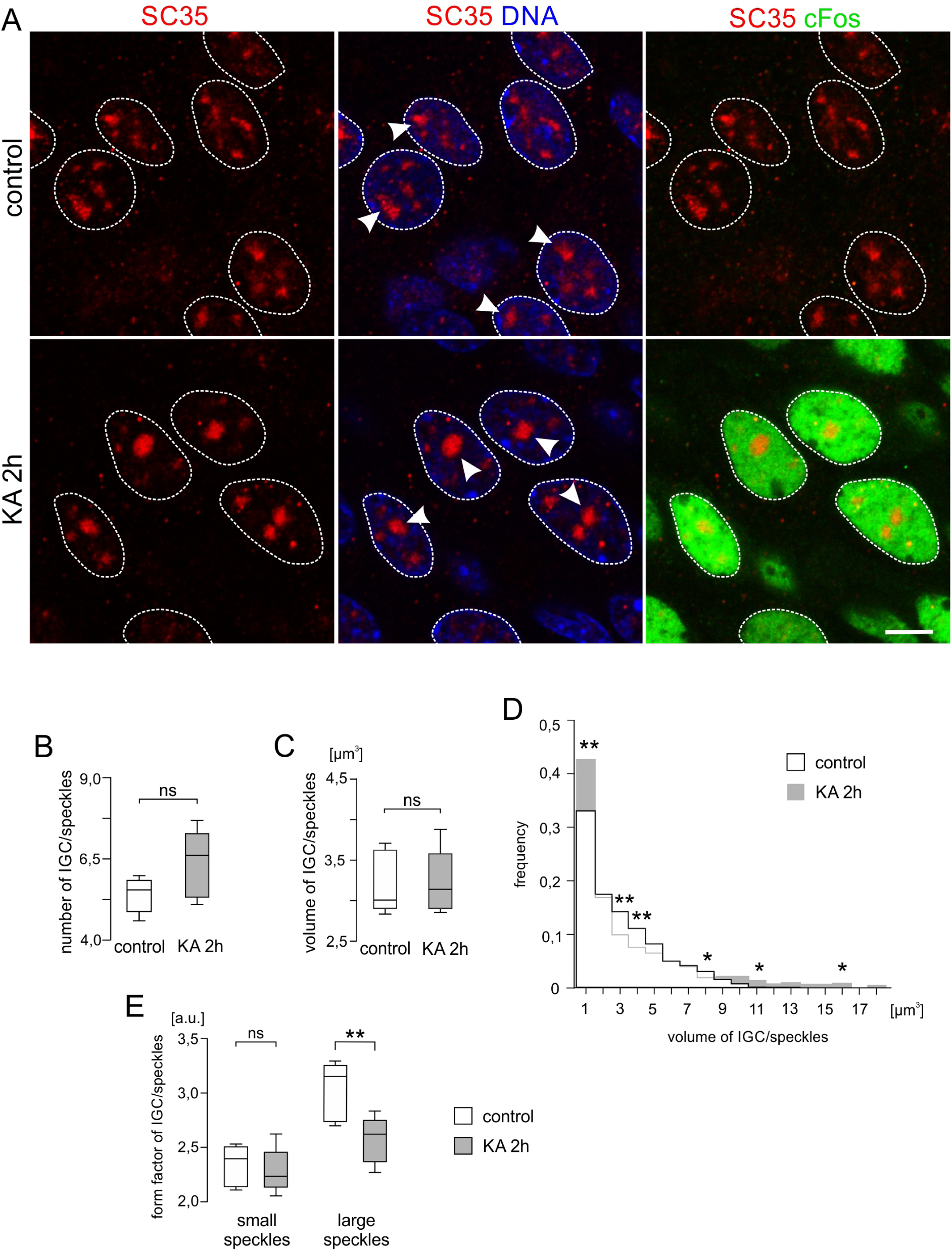
Morphological changes of nuclear speckles upon neuronal activation. **A**. Immunofluorescence labelling of nuclear speckles in the control (control) and 2 h after kainic acid-induced *status epilepticus* (KA 2h). Nuclear speckles were labelled with the SC35 antibody (red). c-Fos protein (green) was used as a marker of neuron activation. DNA was labelled with Hoechst dye (blue). The dotted line indicates nuclear boundaries. Some large nuclear speckles are indicated by the arrowheads. Scale bar: 5 µm. **B**. A mean number of nuclear speckles per granule cell nucleus. **C**. A mean volume of nuclear speckles. **D**. Frequency distribution of nuclear speckles volume. **E**. Form factor for large (>4.75 µm^3^) and small (<4.75 µm^3^) subset of nuclear speckles. Boxes show results for the control (white, control), and for the animals expressing *status epilepticus* (grey, KA 2h). Statistical significance was tested with the Student’s t-test or the Mann-Whitney U test and it is represented by * – p<0.05; ** – p<0.01; ns – not significant. For control N=6 and n=214. For KA 2h N=5 and n=213.

### 3.2. The ultrastructure of the IGC/nuclear speckles is changed upon neuronal activation

To assess changes at the ultrastructural level, we used a serial block-face scanning electron microscopy (SBF-SEM) and the classic transmission electron microscopy (TEM). Although the visualization of IGCs does not require any additional procedures when preparing tissue for canonical imaging in electron microscopy, because they have a characteristic structure, we used a bismuth salts to increase the contrast of ribonucleoproteins (RNPs) (Locke and Huie, 1977) and make IGC/nuclear speckles boundaries detection more easy and certain. This is particularly important in the case of SBF-SEM where enhanced contrast is necessary.

The reconstruction of 3D images from SBF-SEM (Fig. 2A) confirmed the aforementioned morphological changes induced by neuronal stimulation. In the activated neurons of DG large IGC/nuclear speckles were more abundant and more spherical (Fig. 2B and 2C).

**Figure 2.**
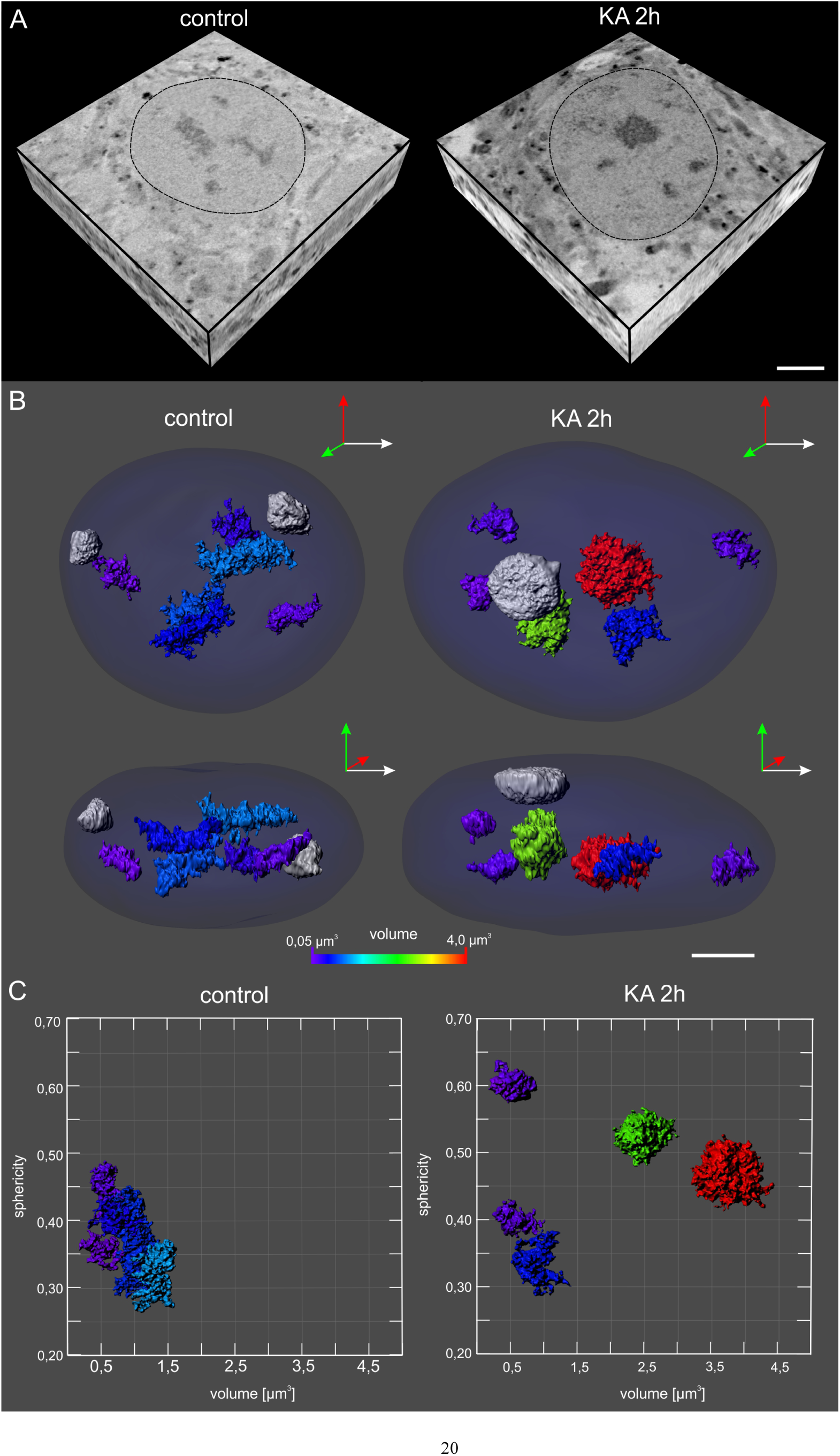
Morphological changes of the IGC/nuclear speckles upon neuronal activation. **A**. Cross-section from a representative 3D stack image for the control (control) and stimulated neuron (KA 2h) obtained using serial block-face scanning electron microscopy (SBF-SEM) by sequential cutting and scanning of a newly exposed surface from a sample resin block. The dashed lines indicate nuclear borders. ICG/nuclear speckles are visible inside the nucleus. The scale bar: 2 µm. **B.** Orthographic projections of the IGC/nuclear speckles 3D reconstruction of in the control (control) and stimulated neurons (KA 2h) obtained using Imaris 7.4.2. from the 3D images shown in A. Nucleoli are depicted in grey. The volume of IGC/nuclear speckles is color-coded according to the bar shown at the bottom. **C.** Plot relating sphericity and volume of the reconstructed IGCs/nuclear speckles from the control (left panel) and stimulated neuron (right panel) from A and B.

On the ultrastructural level, classic TEM analysis showed that control neurons demonstrated typical organization of these nuclear bodies with granules unevenly distributed within the clusters (Fig. 3A, left panel), and the area occupied by them was irregular and hard to define. In the stimulated neurons (KA 2h), the IGC/nuclear speckles were more condensed, with well-distinguishable and smooth peripheries (Fig. 3A, right panel). Moreover, a part of the granules inside the IGC/nuclear speckles strongly aggregated, forming electron-dense compact domains (Fig. 3A right bottom panel), which made up to 15% of the IGC volume (Fig. 3B). The aggregation of granules and formation of the compact domains within IGC/nuclear speckle were visible in more than 60% of stimulated neurons, whereas in control cells they were present in a negligible fraction of cells, in which aggregates encompassed up to 5% of clusters volume (Fig. 3C). In addition, in about 10% of the nuclei of the stimulated neurons, we noticed a separation of the compact domains from the rest of the cluster. Interestingly, after the additional 2 h from the end of status epilepticus, the percentage of nuclei with such ultrastructure increased to 35% (Fig. 4A-4C).

**Figure 3.**
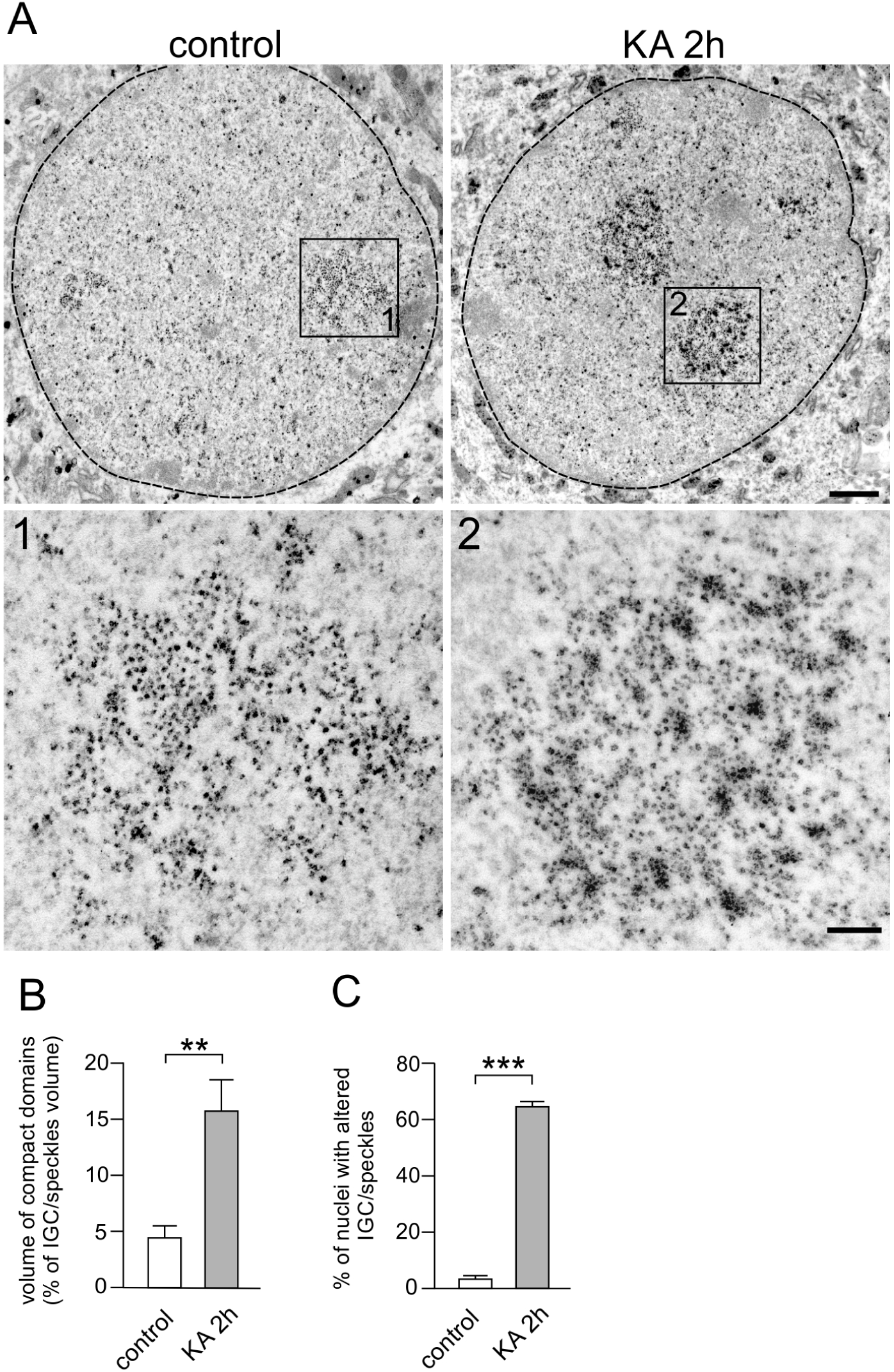
Ultrastructural changes in the IGC/nuclear speckles upon neuronal activation. **A**. Electron micrographs of the granule cell nuclei from the rat dentate gyrus in control (control) and upon *status epilepticus* induced by kainic acid (KA 2h). **1** and **2** show magnified area occupied by IGC/nuclear speckles, marked in the above figures by squares. The dashed, black line indicates nuclear boundaries. Scale bar in the upper panel - 1 µm, in the lower - 250 nm. **B**. Relative IGC/nuclear speckles volume occupied by compact domains. **C**. Percentage of the nuclei containing IGC/nuclear speckles with the compact domains. Bars show results for control animals (white, control), and animals 2 h after kainic acid-induced *status epilepticus* (grey, KA 2h). Statistical significance was tested with the Student’s t-test and it is represented by ** – p<0.01; *** – p<0.001. On B N=5 and n=50 for each of the analysed groups. On C for control N=5 and n=228 and for KA 2h N=5 and n=258.

**Figure 4.**
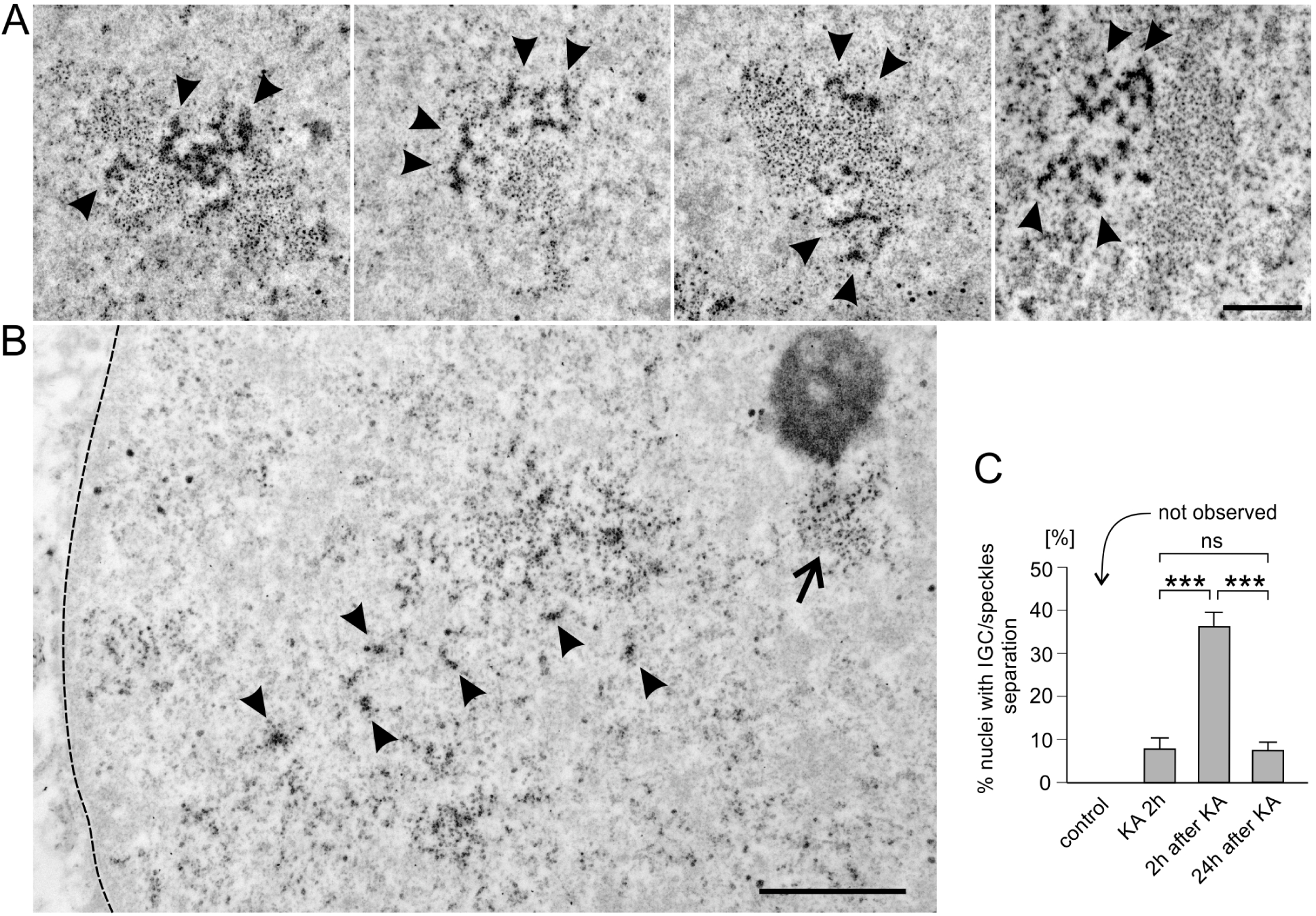
Segregation of compact domains in the IGC/nuclear speckles after the termination of neuronal activation. **A**. IGC/nuclear speckles electron micrographs of rat DG granule neurons 2 h after the completion of the *status epilepticus,* four examples of compact domains (indicated by the arrowheads) separating from non-aggregated IGC domain are shown. Scale bar: 0.75 µm. **B**. Electron micrograph of the rat hippocampal granular neuron nucleus 24 hours after termination of the *status epilepticus*. The arrow shows IGC/nuclear speckle deprived of compact domains, which have been dispersed in the nucleoplasm (indicated by arrowheads). The dashed line shows the nuclear boundary. Scale bar:1 µm. **C**. Percentage of nuclei with compact domains separating from non-aggregated part of IGC/nuclear speckles in the control (control) and treated animals: 2 h after induction KA injection (KA 2h), 2 h (2h after KA) and 24 h (24h after KA) after the end of the *status epilepticus*. Statistical significance was tested with the Student t-test and it is represented by *** – p<0.001; ns - not significant. For each of the analysed groups N=5. For control n=228, for KA 2 h n=258, for 2 h after KA n=75 and for 24 h after KA n=75. Error bars indicate the standard error of the mean.

After 24 h from stimulation, granule aggregates dispersed in the surrounding nucleoplasm (Fig. 4A-4C). Very similar ultrastructural changes in IGC/nuclear speckles were also detected after a short-term stimulation induced by intraperitoneal administration of pentylenetetrazole, an antagonist of GABAA receptors (Fig. S1).

The presence of IGC compact domains in the control samples and the significant increase in their number after stimulation suggest a physiological functionality that so far has not been described in the literature.

The observed changes in the organization of the IGC/nuclear speckles were not associated with neurodegeneration, which we confirmed by Fluoro-Jade C staining (Ikenari et al., 2020) and the TUNEL method (Gavrieli et al., 1992) (data not shown).

**Figure S1.**
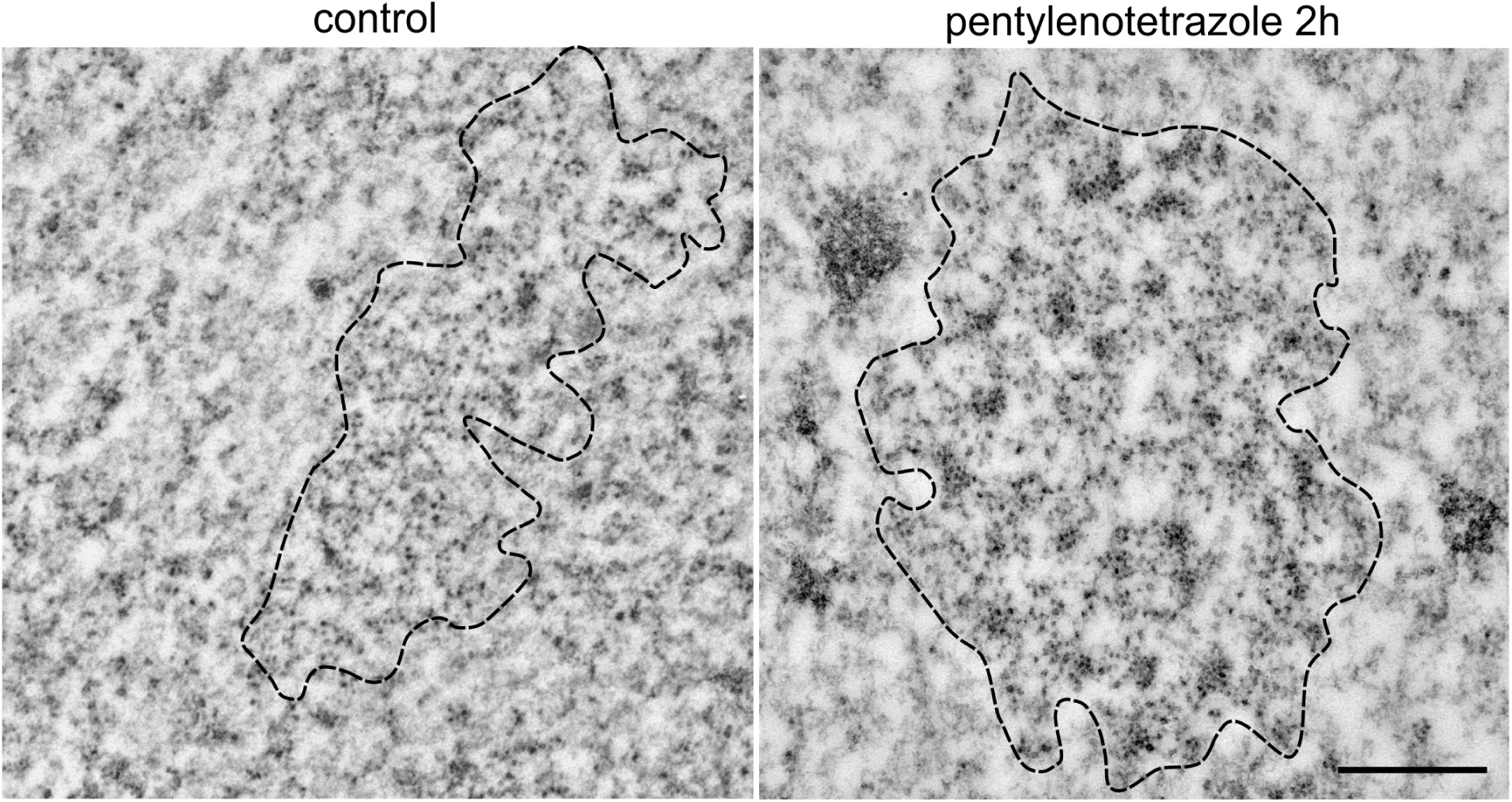
Ultrastructural changes in IGC/nuclear after pentylenetetrazole-induced stimulation. Electron micrographs of IGC/nuclear speckles from the rat’s DG granule cell nucleus in the control and 2 hours after the administration of pentylenetetrazole. In this case contrast of the IGC was not enhanced by bismuth staining. IGC/nuclear speckles are outlined with a dashed line. Scale bar: 0.5 µm.

### 3.3. Stimulation-dependent formation of compact domains within the IGC/nuclear speckles depends on transcription and splicing

To investigate the mechanism underlying the observed changes in the ultrastructure of IGC/nuclear speckles in stimulated neurons, we checked whether they depend on active transcription and splicing. For that purpose, we used an *in vitro* model of organotypic hippocampal cultures stimulated by chemically induced LTP (cLTP) (Otmakhov et al., 2004) to avoid surgical administration of inhibitors directly into the animal hippocampus, Induction of cLTP in CA1 pyramidal neurons caused similar changes in IGC/nuclear speckles ultrastructure as *in vivo* kainate stimulation (Fig. 5).

**Figure 5.**
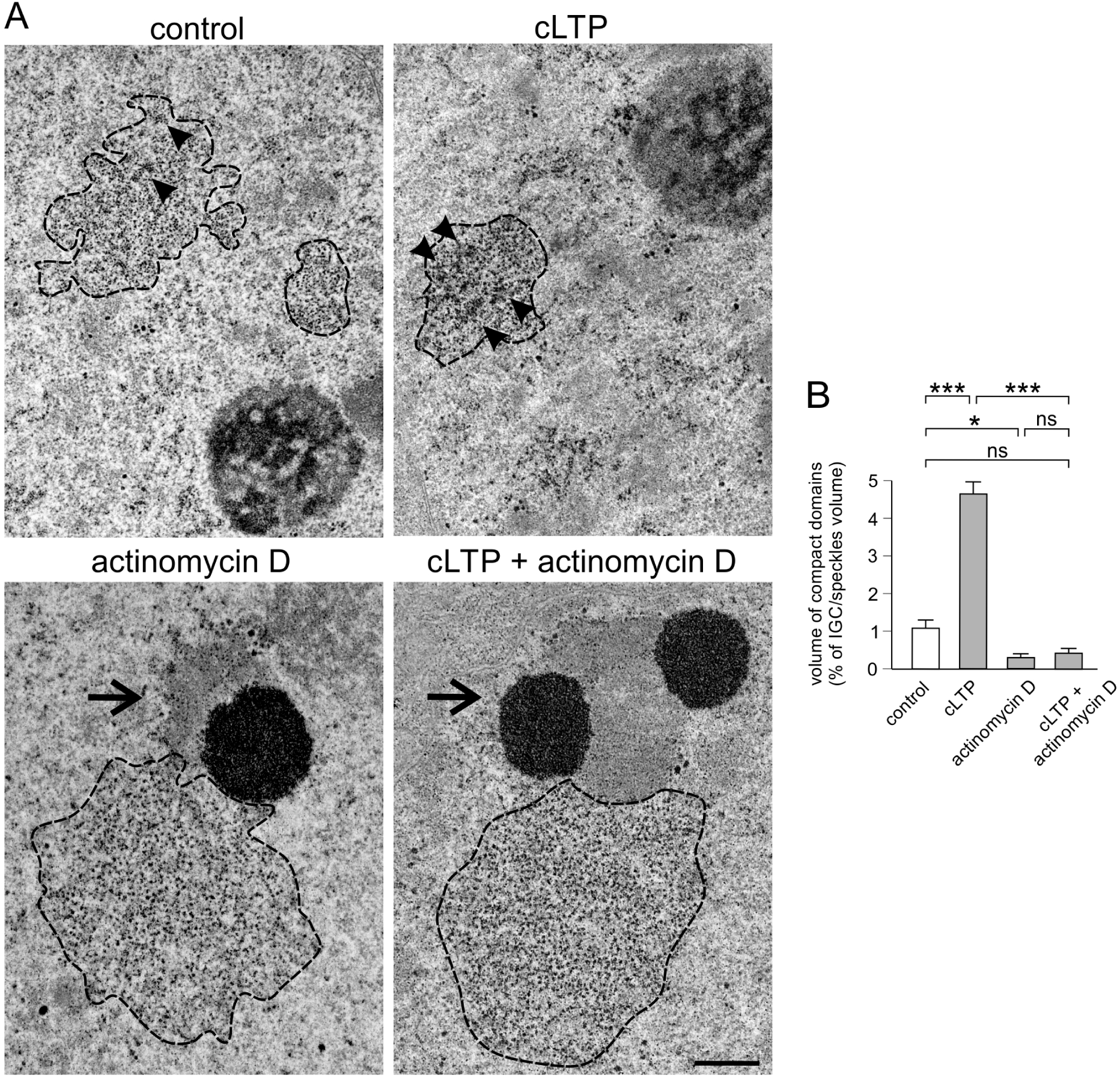
IGC/nuclear speckles in cLTP-stimulated rat hippocampal neurons upon transcriptional inhibition. **A**. Electron micrographs of IGC/speckle in the pyramidal neurons from the organotypic hippocampal slices culture: in the control (control) and chemically evoked LTP (cLTP), and cultures treated with transcription inhibitor (actinomycin D, and cLTP + actinomycin D). The separation of nucleolus components in bottom micrographs is a characteristic feature of transcription inhibition by actinomycin D. The arrowheads indicate compact domains. The dashed line indicates the boundaries of the IGC/nuclear speckles. Segregated nucleoli indicted by the arrows. Scale bar: 0.5 µm. **B**. Relative volume of the IGC/speckle occupied by the compact domains in the cultures treated as in A. Statistical significance was tested with the Student t-test and it is represented by * – p<0.05; *** – p<0.001; ns – not significant. For each of the analysed groups N=5 and n=50. Error bars indicate the standard error of the mean.

To determine the role of active transcription in the structural changes of IGC/nuclear speckles in activated neurons, we blocked transcription using 1 µg/ml concentration of actinomycin D (ActD). As expected, ActD induced separation of the nucleolus dense fibrous and granular components (Reynolds et al., 1964) (Fig. 5). Moreover, it drastically impaired the formation of the compact domains within the IGC/nuclear speckles in stimulated neurons, leading to their complete absence in the majority of nuclear cross-sections (Fig. 5). This result supports the conclusion that the formation of granule aggregates in the IGC/nuclear speckles is a transcription-dependent process. Interestingly, inhibition of transcription induced relocation and aggregation of IGC/nuclear speckles to the proximity of the nucleoli (Fig. 5).

Next, we tested whether the formation of the compact IGC/nuclear speckles domains accompanies pre-mRNA splicing. We used pladienolide B, a splicing inhibitor, which binds to the SF3B complex, a spliceosome component. Pladienolide B induces the formation of defective spliceosomes, which in turn decouples the pre-mRNA splicing (Yokoi et al., 2011). A two-hour incubation of the organotypic hippocampal cultures with this compound significantly increased the number of the IGC/nuclear speckles compact domains. However, the effect appears to be transient as a longer incubation time (8 h) failed to maintain the changes (Fig. 6). This result demonstrates that the short-term formation of compact domains has occurred upon splicing inhibition.

**Figure 6.**
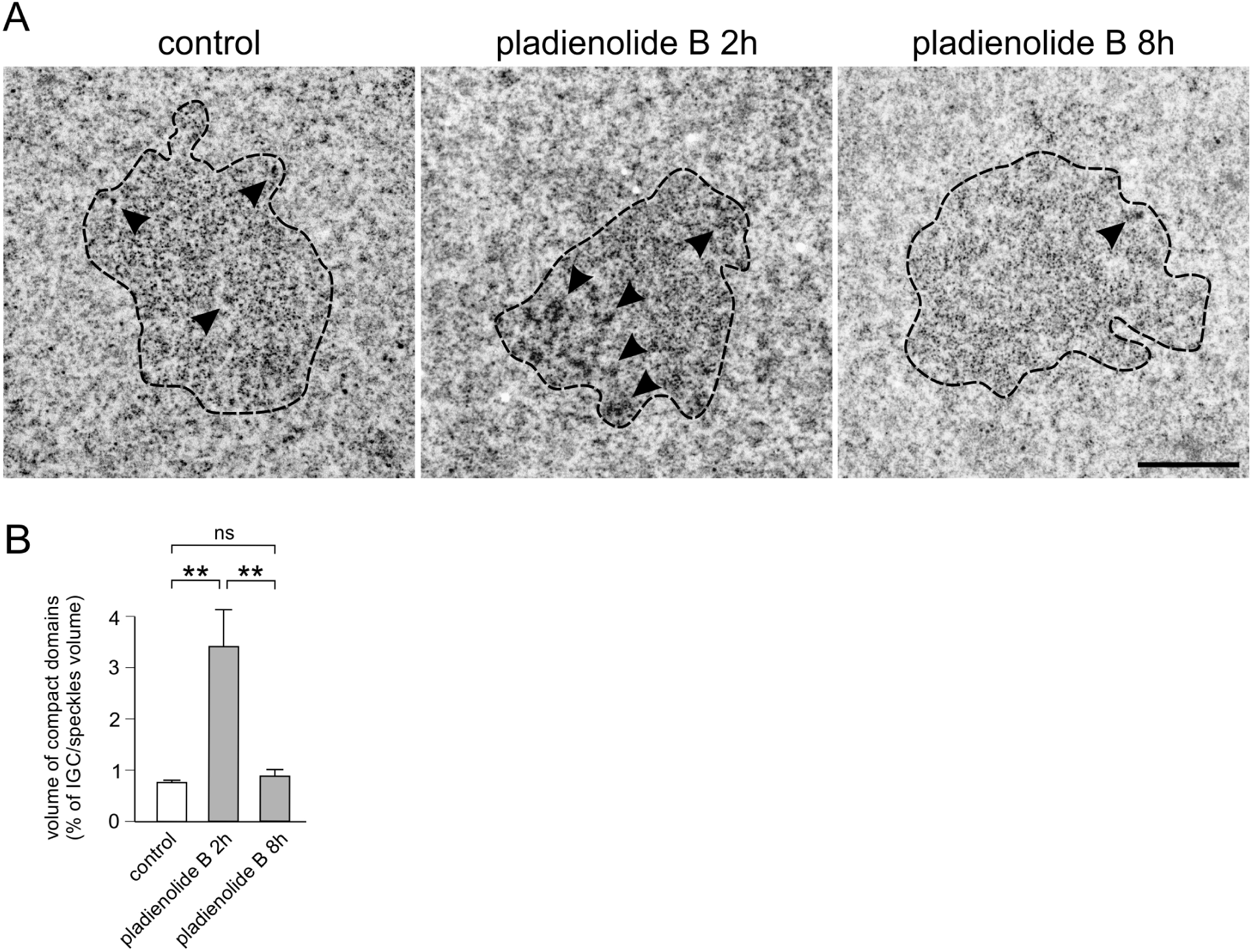
IGC/nuclear speckles after inhibition of mRNA splicing. **A.** Electron micrographs of the IGC/nuclear speckles in the pyramidal cells from rat’s hippocampal organotypic culture in control (control), 2 hours (pladienolide B 2h), and 8 hours (pladienolide B 8h) after mRNA splicing inhibition. The arrowheads indicate compact domains. The dashed line indicates the boundaries of the IGC/nuclear speckles. Scale bar: 1 µm. **B**. Relative volume occupied by compact domains from cultures treated as in A. Statistical significance was tested with the Student t-test and it is represented by ** – p<0.01, ns – not significant. For each of the analysed groups N=5 and n=50. Error bars indicate the standard error of the mean.

### 3.4. Neuronal activation induces changes in the localization of IGC/nuclear speckles proteins

A formation of the compact IGC/nuclear speckles domains, observed in neurons after stimulation, may be related to a change in their protein composition and/or protein quantity.

Therefore, we performed an immunogold labelling of the selected IGC/speckle proteins in kainate-activated hippocampal granular DG neurons. As the number of gold particles within the IGC/nuclear speckles is linearly proportional to the antigen accumulation (Amiry-Moghaddam and Ottersen, 2013), immunogold labelling allows for both qualitative and quantitative analysis.

The SR proteins, mRNA splice factors, are characteristic and important components of nuclear speckles. To determine the localization of such proteins, we used the antibody which recognizes phosphorylated epitopes of SR proteins (Turner and Davies, 1986) and has been previously used for their immunolocalization at the ultrastructural level (Niedojadlo et al., 2012; Tripathi et al., 2012). In the DG neuronal nuclei from the stimulated animals, we observed an almost twofold increase in the labelling density of phosphorylated SR proteins within the IGC/nuclear speckles (Fig. 7, shown in the upper panel). Moreover, labelling was stronger at the periphery of the compact domains (Fig. 7, shown in the upper panel).

**Figure 7.**
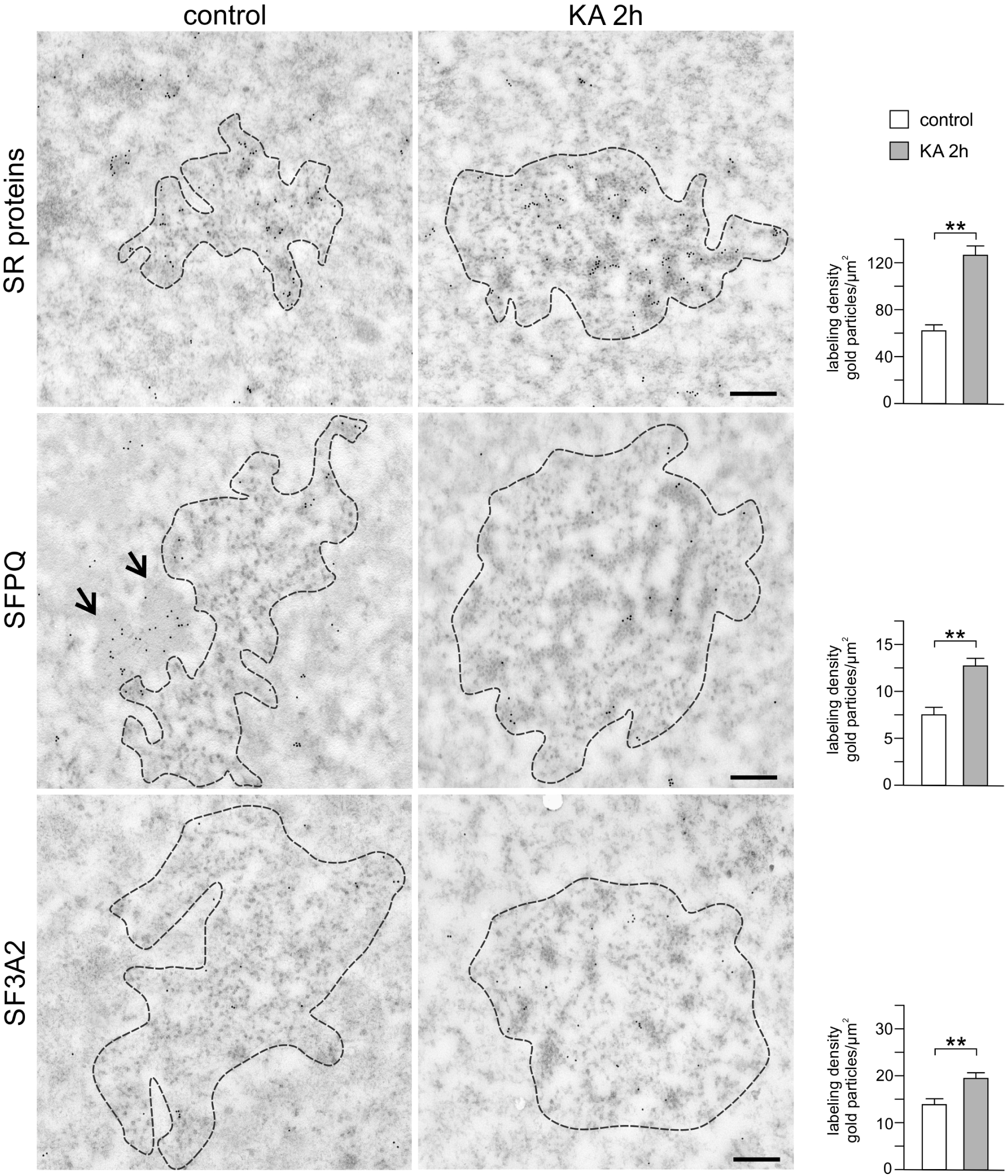
Localization of IGC/nuclear speckle components upon neuronal activation. Immunogold labelling with an antibody against SR, SFPQ, and SF3A2 proteins. Electron micrographs of IGC/nuclear speckles in granular cells from rat’s DG in the control (control) and 2 hours after the induction of *status epilepticus* by kainic acid (KA 2h). The arrows show IGAZ for which the SFPQ protein is a characteristic component. The dashed line indicates the IGC/speckle boundaries. The scale bar: 0.25 µm. Graphs represent labelling density in IGC/nuclear speckles profiles in the control (control, white) and 2 hours after the induction of *status epilepticus* by kainic acid (KA 2h, grey). Statistical significance was tested with the Student t-test and it is represented by ** – p<0.01. For each of the analysed groups N=3 and n=30. Error bars indicate the standard error of the mean.

The SFPQ (*Splicing factor, proline- and glutamine-rich*), also known as PSF (*PTB-associated-splicing factor*, *Polypyrimidine tract-binding protein-associated-splicing factor*) is a multifunctional RNA-binding nuclear protein. SFPQ is involved in several stages of mRNA maturation and, among others, is a splicing factor essential for the formation of the spliceosome and stage II catalysis of the pre-mRNA splicing (Yarosh et al., 2015). It is also a protein characteristic of the paraspeckles, another nuclear body, often present in the vicinity of IGC/nuclear speckles. Similar to nuclear speckles, paraspeckles have an ultrastructural equivalent, called IGAZ (*Interchromatin Granule-Associated Zones*), but it is not clear whether they correspond entirely (Fox and Lamond, 2010). The presence of SFPQ has also been reported within IGC/nuclear speckles of hepatocytes (Saitoh et al., 2004). In line with these observations, immunogold labelling of the control sections revealed the presence of SFPQ within the IGC/nuclear speckles from resting DG neurons. After neuronal stimulation, the labelling density of the SFPQ epitopes in these structures almost doubled (Fig. 7, shown in the middle panel). The SFPQ-corresponding particles were predominantly located within the IGC/speckle compact domain or at the boundary of this domain. As expected, the SFPQ labelling density in IGAZ/paraspeckles was much higher than in the IGC/nuclear speckles (see the control of Fig. 7, middle panel).

Another IGC/speckle protein, SF3A2 (*splicing factor 3A subunit 2*, other abbreviations used: SF3a66 or SAP62) belonging to the SF3A subcomplex of the U2 snRNP, is a component of the spliceosome and a commonly used marker of nuclear speckles (Will and Luhrmann, 2011). After stimulation, a level of SF3A2 protein increased slightly in IGC/nuclear speckles (Fig. 7, the lowest panel) and the protein localized in both IGC/nuclear speckles compartments – in compact domains (mainly at their edges) and in part where granules are more dispersed.

Since RNA polymerase II (RNAPII) was found in the nuclear speckles of non-neuronal cells (Bregman et al., 1995; Saitoh et al., 2004), we decided to check whether it is also present in these structures in the neuronal cells and whether the neuronal activity, which significantly stimulates transcription of hundreds of genes, has an impact on the concentration and localization of RNAPII epitopes in the IGC/nuclear speckles. We chose two forms of activated RNAPII for labelling, phosphorylated on Serine 5 (S5) and on Serine 2 (S2) of its co-catalytical carboxyterminal domain (CTD), which represents the active form of the protein. For the S5 phosphorylated form of this polymerase, the labelling density after stimulation is about one-third higher than in the control (Fig. 8, top panel), whereas the S2 phosphorylated form density under the same conditions is even five times higher than in the control (Fig. 8, bottom panel). The above-described result shows that both activated forms of RNAPII are present in the neuronal IGC/nuclear speckles and their levels are increased by neuronal stimulation.

**Figure 8.**
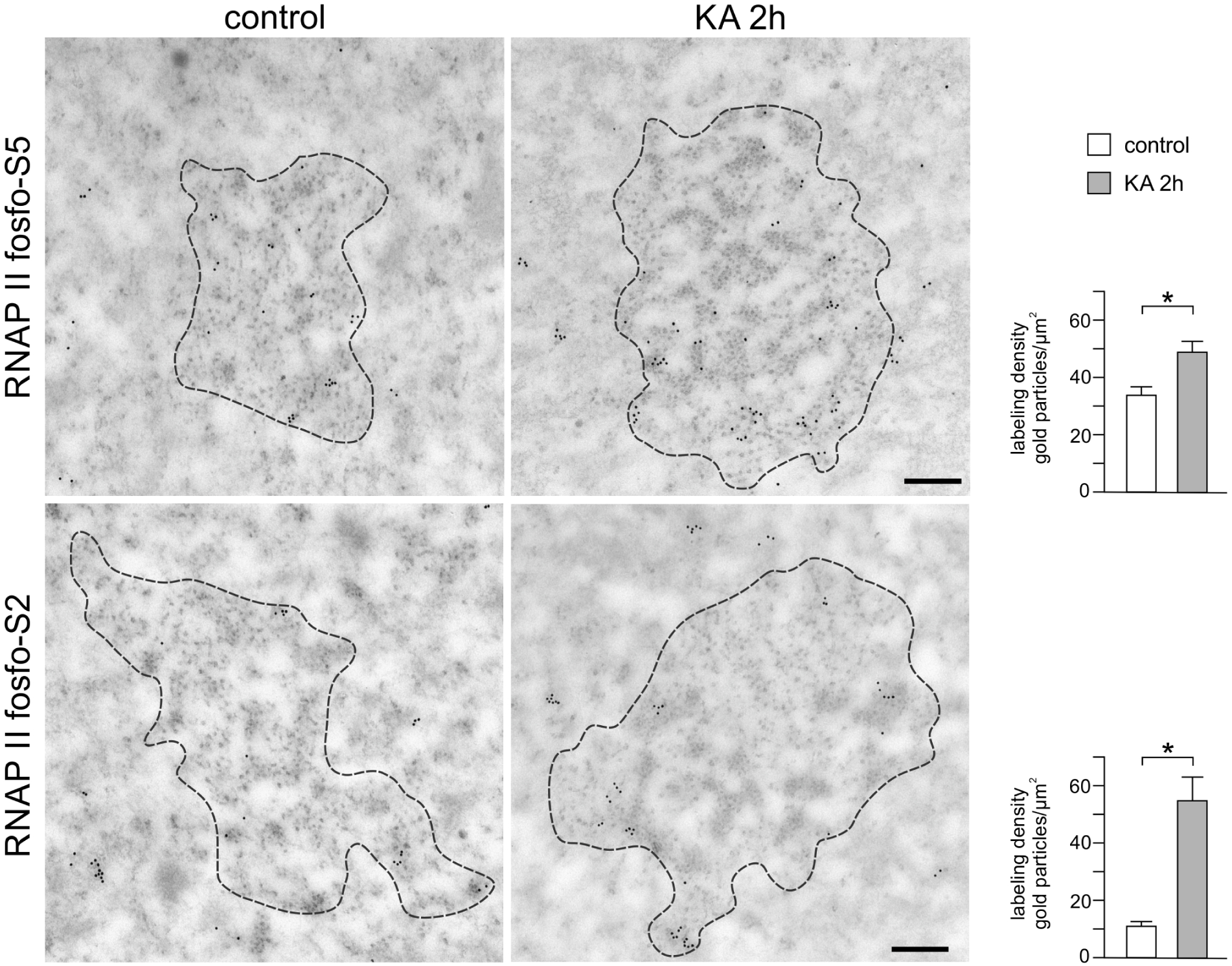
Localization of RNAP II in IGC/nuclear speckles upon neuronal activation. Electron micrographs of IGC/nuclear speckles in granular cells from rat DG, in control (control) and 2 h after the induction of *status epilepticus* by kainic acid (KA 2h) showing immunogold labelling with an antibody recognizing activated forms of RNA polymerase II (RNAP II phospho-S5, and RNAP II phospho-S2 epitopes). The dashed line indicates IGC/nuclear speckles boundaries. The scale bar: 0.25 µm. Graphs represent labelling density in IGC/nuclear speckles profiles in control (control, white) and 2 h after the induction of *status epilepticus* by kainic acid (KA 2h, grey). Statistical significance was tested with the Student t-test and it is represented by * – p<0.05. For each of the analysed groups N=3 and n=30. Error bars indicate the standard error of the mean.

Finally, to confirm the quality of our immunogold staining, as a negative control, we used an antibody against coilin, a protein characteristic for the Cajal bodies (Morris, 2008), which are often observed in the vicinity of IGCs/nuclear speckles (Lafarga et al., 1998; Lafarga et al., 2017). The immunogold labelling revealed that coilin was specific for the Cajal bodies and was not detected in the IGCs/nuclear speckles under control and stimulating conditions (Fig. S2).

**Figure S2.**
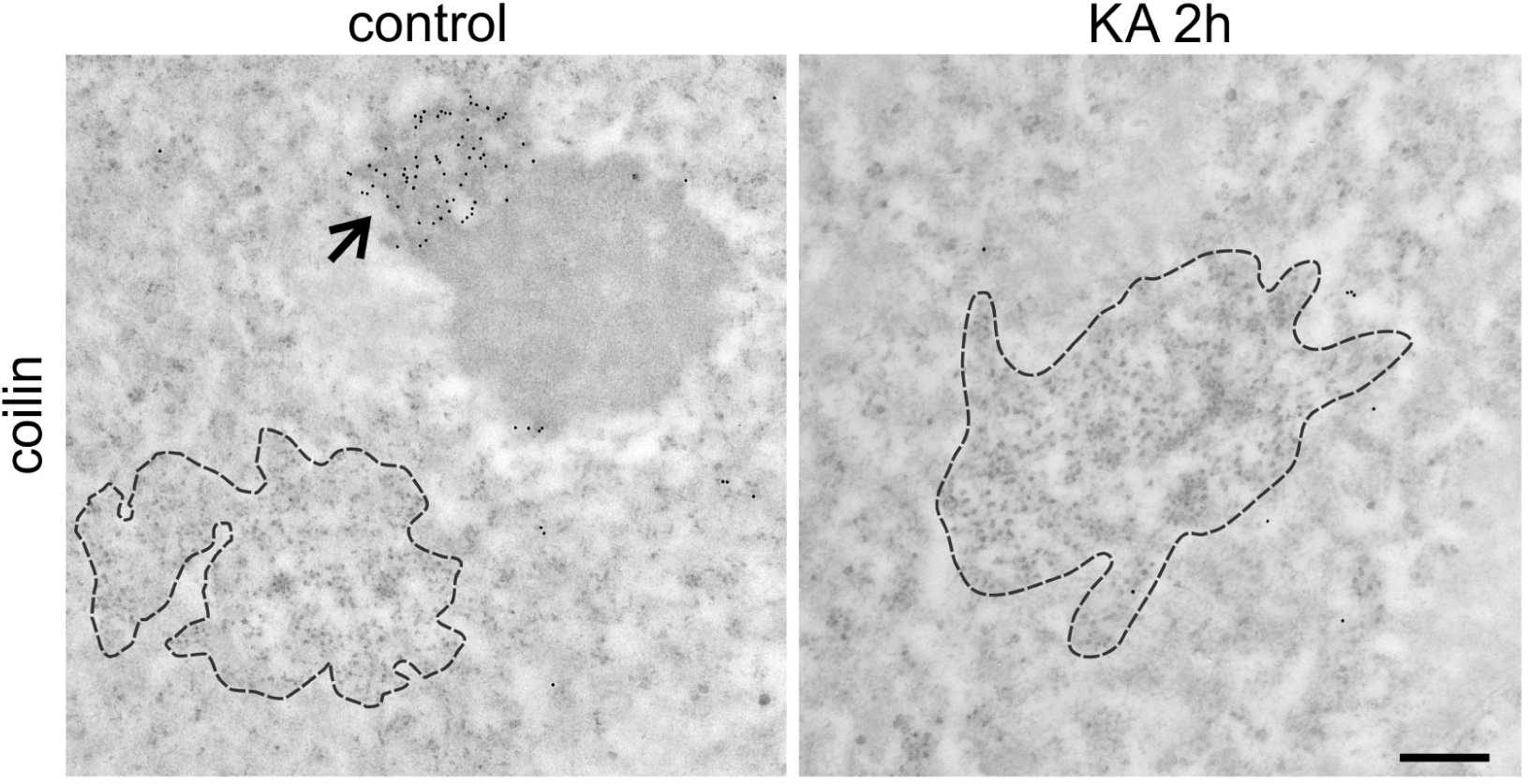
Coilin is not localized in the IGC/nuclear speckles upon neuronal activation. Immunogold labelling with an antibody against coilin, a characteristic component of Cajal bodies. Electron micrographs of IGC/nuclear speckles in granular cells from the rat DG, in the control (control), and 2 hours after the induction of *status epilepticus* by kainic acid (KA 2h). The arrow in the control micrograph shows the Cajal body adjacent to the nucleolus. The dashed line indicates the IGC/speckle boundaries. The scale bar: 0.25 µm.

### 3.5. DNA localization in IGC/nuclear speckles

As it was mentioned above, numerous studies reported the presence of the specific gene *loci* within the nuclear speckles or at their periphery. Therefore, we decided to check whether the formation of the compact domains in activated neurons is accompanied by the presence of DNA. To determine the location of DNA within the compact domains, we performed labelling of the DNA ends on the ultrathin section as cutting the resin-embedded tissue sections with an ultramicrotome produces free DNA ends (Thiry, 1992). Such produced ends can be tagged with fluorescein-labelled nucleosides using TdT polymerase and then detected with anti-fluorescein antibody conjugated with colloidal gold. Consequently, DNA labelling showed its high density in the heterochromatin, while it significantly lowered in the IGC/nuclear speckles interior (Fig. 9). In contrast, we found no statistically significant difference in the labelling density of the IGCs/nuclear speckles under control and stimulating conditions. However, in the stimulated neurons DNA labelling was more evident in the interior of IGC/nuclear speckles than in the proximity of their boundary (Fig. 9C).

**Figure 9.**
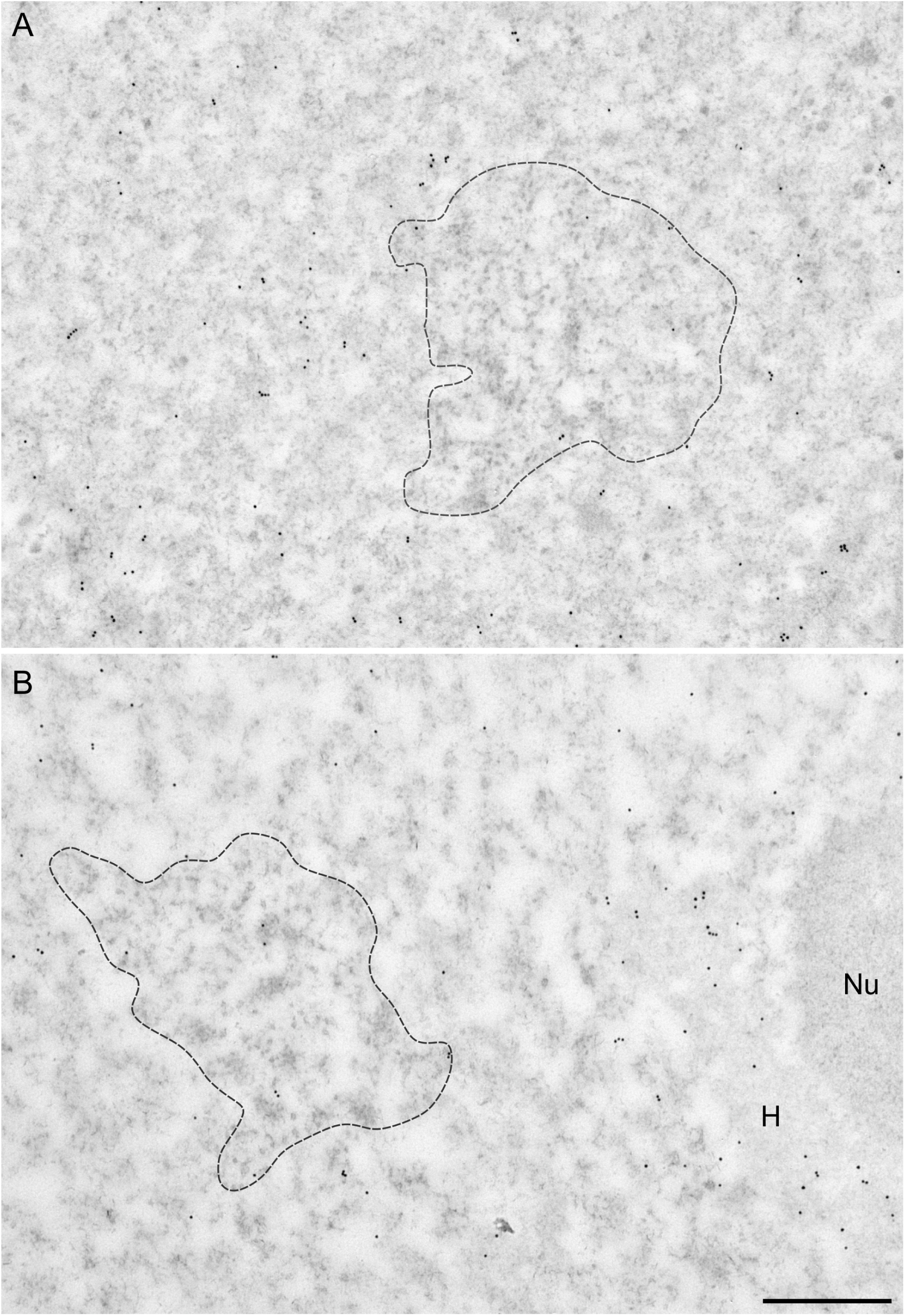
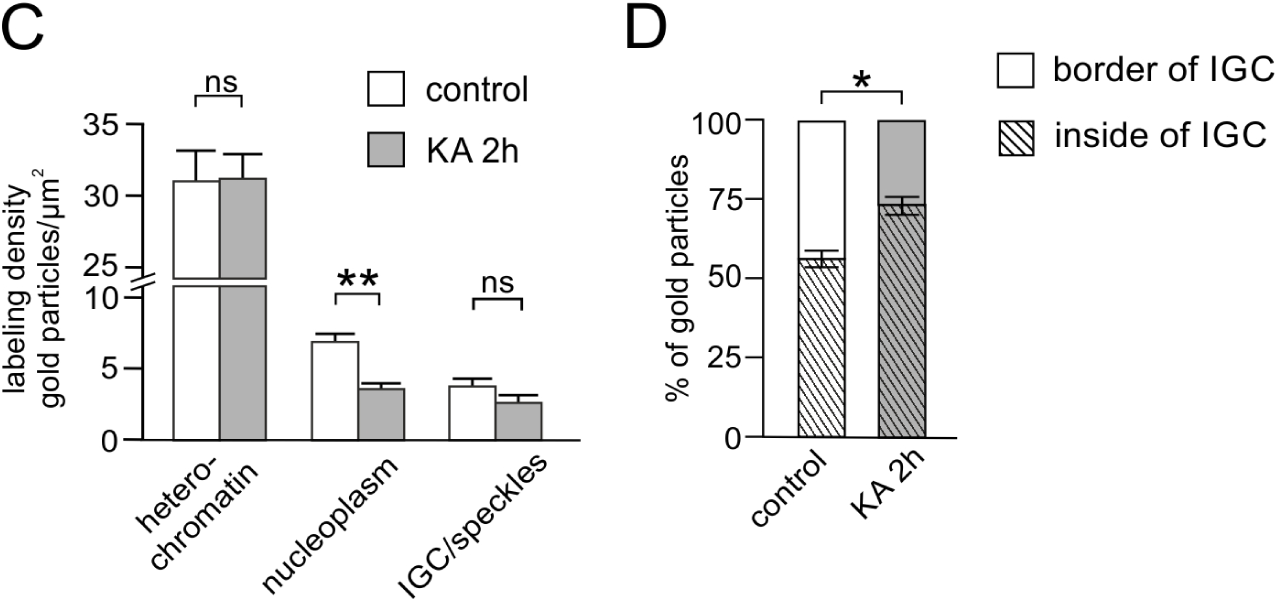
DNA localization in IGC/nuclear speckles upon neuronal activation. Immunogold labelling of fluorescein-tagged nucleotides. Electron micrographs of IGC/nuclear speckles in granular cells from rat DG, in control (**A**), and 2 hours after the induction of *status epilepticus* by kainic acid (**B**). The dashed line denotes the IGC/nuclear speckles borders, H – heterochromatin, and Nu – nucleolus. Scale bar: 0.5 µm. **C**. Labelling density of DNA in heterochromatin, nucleoplasm, and IGC/nuclear speckles. **D**. Percentage of gold particles marking DNA localization at the edge of the nuclear speckles and inside the IGC/nuclear speckles. Bars show results for control animals (white, control), and animals 2 hours after kainic acid-induced *status epilepticus* (grey, KA 2h). Statistical significance was tested with the Student’s t-test and it is represented by * – p<0.05; ** – p<0.01; ns – not significant. For each of the analysed groups N=5 and n=50. Error bars indicate the standard error of the mean.

### 3.6. Proteomic, functional, and quantitative analysis of the proteins identified in neuronal nuclear fraction enriched in nuclear speckles

Significant changes in the organization of neuronal nuclear speckles and the differences in the levels of some nuclear speckles proteins suggest that neuronal stimulation may lead to the modification of their proteome. In line with such a possibility, we decided to verify this by qualitative and quantitative mass spectrometry analysis of the nuclear speckles enriched nuclear fraction (Mintz et al., 1999), isolated from the hippocampi of control and *status epilepticus*-evoked animals. A fractionation efficiency was confirmed by the western blot analysis of the SC35 (SRSF2) protein (Fig. S3).

**Figure S3.**
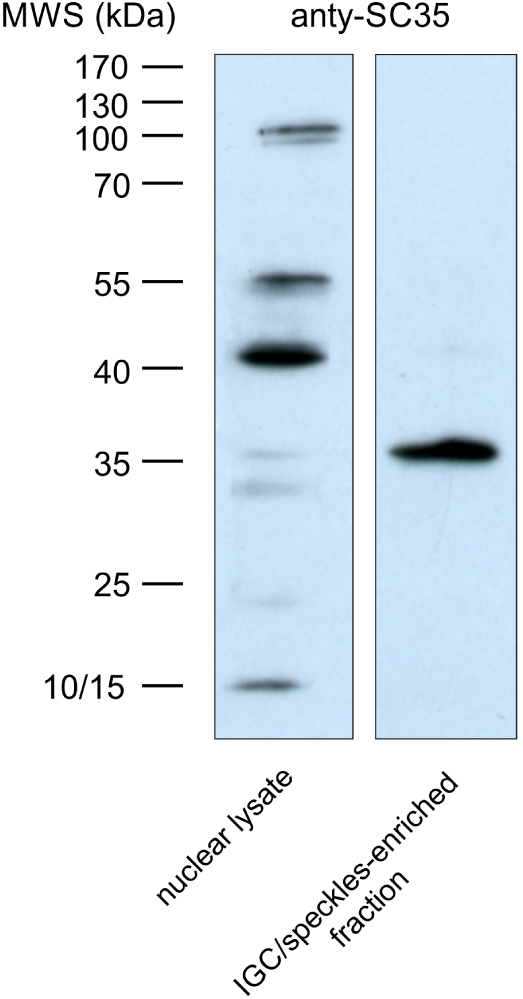
Western blot analysis of the nuclear fraction enriched in nuclear speckles. The left lane presents a nuclear lysate isolated from the rat hippocampus, and the right one, lysate from the nuclear speckles-enriched fraction. In both lanes, 4 μg of total protein was run, and an anti-SFRS2 (SC35) antibody was used for detection. Both lanes originate from the same nitrocellulose membrane. MWS, molecular weight standard.

By comparing a list of the peptides identified by the MS in the nuclear speckles-enriched fraction against the UniProt and Gene Ontology (GO) database, we identified 887 proteins of which 491 were nuclear. The remaining proteins - most probably representing the contamination - were non-nuclear proteins (Tab S4 in supplementary Excel file), except for the three cytoskeletal proteins (two cytoplasmic actins and one unconventional myosin) which, are known to be present in the nuclei, and have been found in the nuclear speckles (Naum-Ongania et al., 2013; Pranchevicius et al., 2008; Saitoh et al., 2004).

Subsequently, similarly to Saitoh (Saitoh et al., 2004), we divided the identified nuclear proteins into three groups: (1) known speckle proteins (described previously by Galganski in (Galganski et al., 2017)) or assigned as speckle proteins by GO analysis, altogether 119 proteins (shown in red in Tab. S2 of supplementary Excel file); (2) potential speckle proteins with the function or structure similar to the already known speckle components or belonging to the protein families known to be present in nuclear speckles, 106 proteins (shown in green in Tab. S2 of supplementary Excel file); and finally the (3) known nucleoplasmic proteins, which were most likely contamination, 266 proteins (shown in Tab. S3 of supplementary Excel file).

Further, we used the UniProt database to determine, which of the identified proteins contains RNA-binding domains (RMM, KH, and DRBM domains) as well as the SMART database (Letunic and Bork, 2018) to identify LCR regions, characteristic for proteins found in the nuclear speckles (Saitoh et al., 2004; van der Lee et al., 2014) (Tab. S2 of supplementary Excel file).

Almost 60% of the proteins from the first group (shown in red in Fig. 10 and Tab. S2 of the supplementary Excel file) were related to mRNA metabolism (71/119), and most of them were pre-mRNA splicing factors (52%, 63/119) or proteins related to mRNA export (8%, 10/119). The second functional group, constituting 30% of the identified known speckle proteins were transcription factors (35/119). In addition, several proteins were related as chaperones, proteins associated with chromatin organization and/or histone modification, DNA repair, protein degradation, signal transduction, or described by the UniProt database as multifunctional factors. Among 106 proteins identified as potential components of nuclear speckles (shown in green in Fig. 10, and Tab. S2 of supplementary Excel file), the most abundant group, 44%, (47/106) includes factors functionally associated with transcription. Functions related to mRNA metabolism were again attributed to a high fraction (26%, 28/106) of the identified proteins. This includes pre-mRNA splicing factors, constituting 20% of the total hits (22/106). Similarly to the first group, the potential speckle proteins also included proteins involved in the organization of chromatin and modification of histones, protein degradation, signal transduction, DNA repair, chaperones, and multifunctional factors. Summarizing, the majority of speckle proteins identified in neurons were involved in pre-mRNA splicing and maturation, functions previously described for non-neuronal cells (Galganski et al., 2017; Saitoh et al., 2004). However, the second largest group found in our analysis were proteins involved in the regulation of transcription-a feature presumably specific to the neuronal cells.

**Figure 10.**
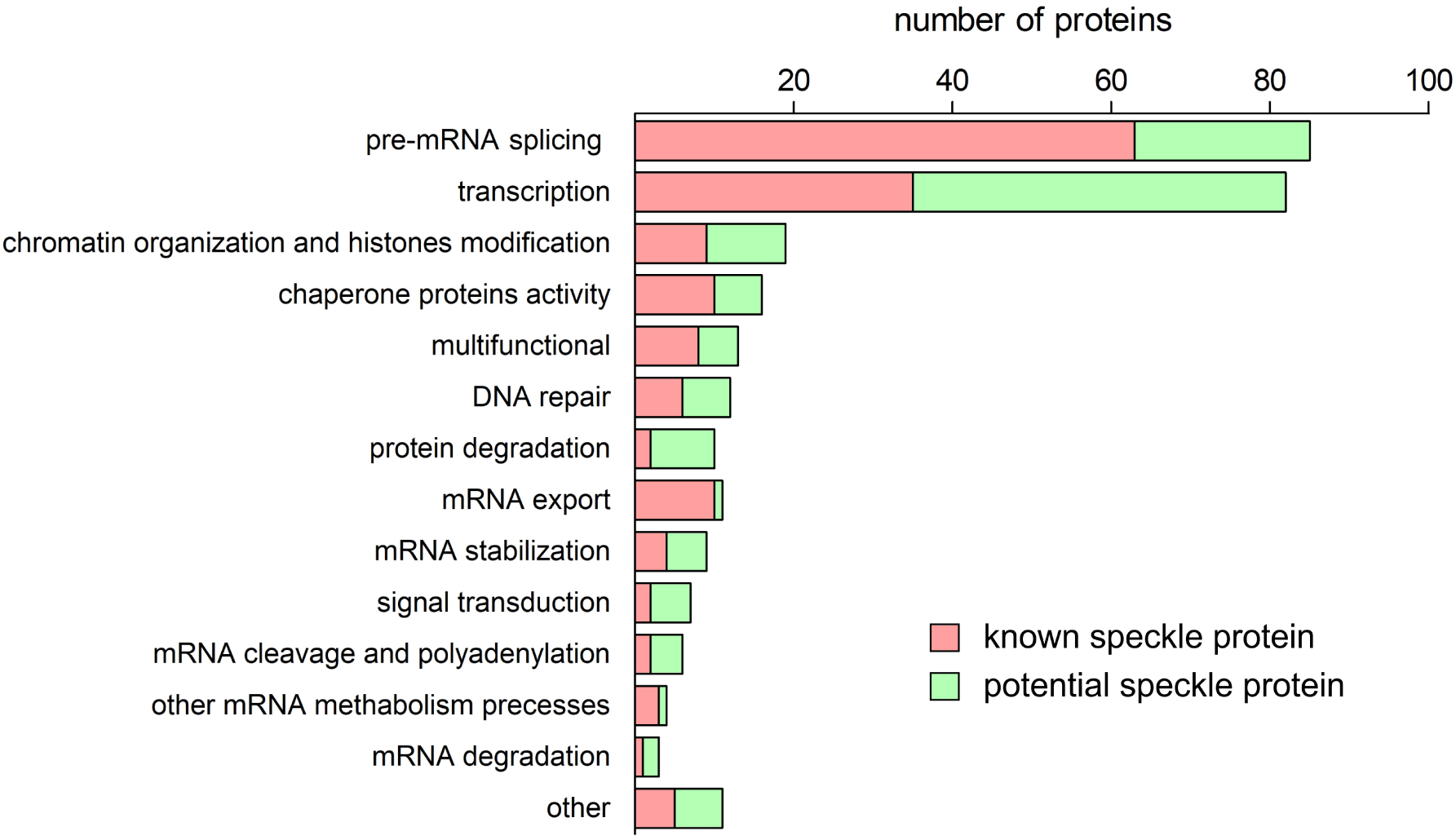
Functional analysis of proteins identified in the nuclear speckles-enriched fraction from neuronal cells. Protein functions were determined using the UniProt database. Proteins with more than one function were assigned into several functional categories. Known speckle proteins are highlighted in red and potential speckle proteins in green. The scale shows the number of identified proteins.

Finally, we performed a quantitative analysis comparing the number of peptides detected for the individual proteins, in the control and kainate-stimulated animals (Tab. S2 of supplementary Excel file). Neuronal stimulation increased the level of 29 proteins and led to a decrease in the level of only two proteins (Fig. 11, and Tab. S2 of supplementary Excel file). Almost 1/3 of the up-regulated proteins (9/29) belong to hnRNPs functional group: A1, A2/B1, A3, H, U, H2, D-like, C, and L. Seventeen proteins (17/29) are involved in splicing (e.g. U5 small nuclear ribonucleoprotein, 200 kDa helicase, YTH domain-containing protein 1, splicing factor 3A subunit 2 and almost all hnRNPs), and 13 (13/29) are transcriptional factors (e.g. scaffold attachment factor B1, metastasis-associated protein MTA1, transcription activator BRG1, hnRNP D-like and some others hnRNPs). Other functionalities, such as chromatin re-organization, DNA repair, or mRNA export were represented rarely (Fig. 11, and Tab. S2 of supplementary Excel file). In contrast, our analysis revealed that only two proteins were downregulated: specifically, cytoskeletal actin isoforms 1 (β-actin) and 2 (γ-actin), both of which were previously identified within nuclear speckles (Naum-Ongania et al., 2013; Saitoh et al., 2004).

**Figure 11.**
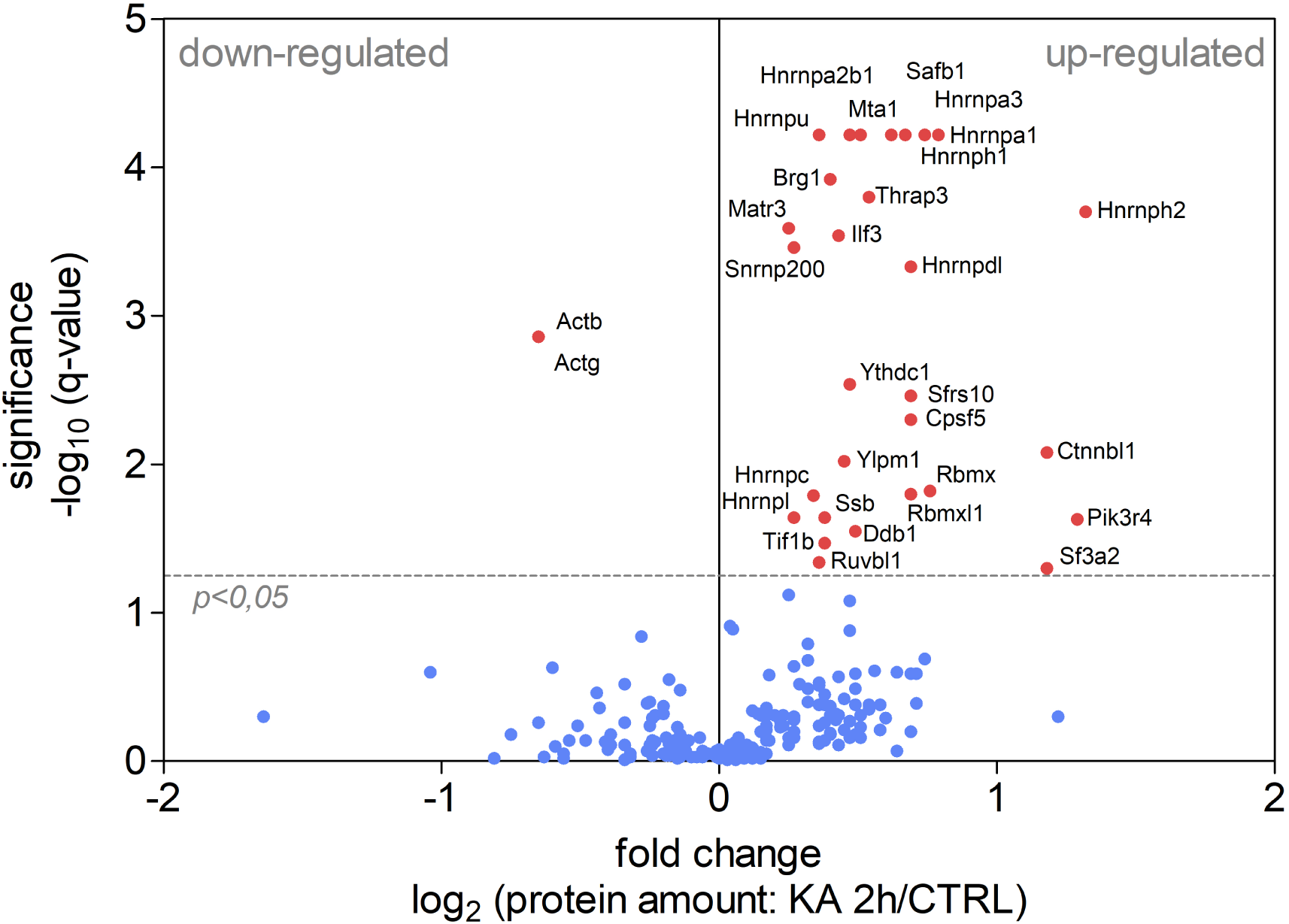
Quantitative analysis of the changes in proteome in the nuclear speckles-enriched fraction from neuronal cells in response to activity. The volcano plot of mass spectrometry results comparing the level of proteins identified in the nuclear speckles-enriched fraction from the control animals (CTRL) and 2 hours after the induction of *status epilepticus* by kainic acid (KA 2h). Each point represents one protein; the level of statistical significance is shown on the Y axis, and the log2 fold change is shown on the X axis. Proteins with levels significantly changed are marked in red. The complete data set is presented in Table S2.

## 4. Discussion

### 4.1. Neuronal activation-induced changes in morphology of the IGC/nuclear speckles that are dependent on transcription and splicing

The convulsion stimulation of hippocampal formation, results in a transient increase of transcription of many genes, including early-response genes (Fernandez-Albert et al., 2019). This is accompanied by covalent modifications of histone tails (Huang et al., 2002; Sawicka and Seiser, 2012; Taniura et al., 2006), chromatin structure rearrangement (Grabowska et al., 2022; Tao-Cheng, 2018), displacement of certain gene *loci* (Walczak et al., 2013), and reorganization of the PML bodies (Hall et al., 2016). The results presented in this study demonstrate that experimental neuronal stimulation has a significant influence on the IGC/nuclear speckle organization and protein composition. In the stimulated granular neurons, nuclear speckles became oblong with more defined borders. This is accompanied by the formation of internal clusters (aggregates) of speckle granules, which we refer to as compact domains. Such compact domains are present both *in vivo* and *in vitro*. In stimulated cells, with the same average number of nuclear speckles per nucleus, we observed a significantly higher number of small and a slightly higher number of large nuclear speckles, exceeding the volumes of those present in the control. This may suggest that the nuclear speckles undergo the process of speckle fission and/or fusion. We do not exclude the possibility of mobility of the nuclear speckles and their association as it was previously observed in the CHO cells after blocking the transcription (Kim et al., 2019). Structures similar to compact domains have been previously observed in various experimental setups. Fedorko and Hirsch noted their presence in fibroblasts after treatment with mepacrine, an antiprotozoal drug (Fedorko and Hirsch, 1969). Also, Miyai and Steiner showed similar structures in the hepatocytes of rats fed with food containing the carcinogen, ethionine (Miyai and Steiner, 1965). Furthermore, Svoboda and Higginson (Svoboda and Higginson, 1968), as well as Krzyżowska-Gruca et al. observed such compact domains in rat hepatocytes after intraperitoneal administration of hepatocarcinogens and the cytostatic, nitracrine (Krzyzowska-Gruca et al., 1983). However, while given examples relate to pathological conditions, the presence of the compact domains in the control neurons indicates their physiological role, not related to any dysfunction. This suggestion seems to be confirmed by Pena and co-authors who, under normal conditions, observed electron-dense aggregates in the IGC/nuclear speckles of the sensory neurons from the rat trigeminal ganglion (Pena et al., 2001). Similar structures were also described by Jones and LaVelle in the Chinese hamster facial nerve maturation (Jones and LaVelle, 1986). None of the studies explored the role or made any speculation regarding the nature of these structures. Interestingly, similar compact domains were observed by Huang in tsBN2 fibroblasts lacking functional RCC1 (Regulator Of Chromosome Condensation 1), showing an impaired export of poly(A)+ RNA to the cytoplasm. The accumulation of pre-mRNA splicing factors and poly(A)+ RNAs in the nuclear speckles induced morphological changes such as size increase and perimeter smoothening, and on the ultrastructural level, the formation of aggregates, observed also in the nucleoplasm. Importantly, the aggregates observed by Huang were resistant to the RNase A and DNase I treatment and contained SC35 (SRSF2) protein (Huang et al., 1997).

The inhibition of transcription in neurons resulted in the complete absence of compact domains in IGC/nuclear speckles upon stimulation. The effect was very strong as the number of compact domains was not only lower than in stimulated neurons without inhibited transcription but also lower than in the control, untreated neurons. Additionally, we observed an increase in the nuclear speckles volume, their rounding up, and relocation to the proximity of the nucleolus. The latter effects of the inhibition of transcription with ActD were also described by Schoefl in the early research on the ultrastructure of nucleoli originating from *Guinea baboon* kidney cell cultures (Schoefl, 1964). Contrary to these observations, in non-neuronal cells, blocking of transcriptional activity seems to induce the presence of compact domains in IGC/nuclear speckles. Schoefl visualized aggregates after 7 h-long incubation with ActD (Schoefl, 1964). Similar compact domains were visible in IGC/nuclear speckles of mouse hepatocytes, after 9 h of α-amanitin administration (Marinozzi and Fiume, 1971). A more recent report by Spector et al. (Spector et al., 1993) showed small aggregates in the IGC/nuclear speckles from the CHO cells after 5-h inhibition of transcription with α-amanitin. Overall, it seems to be most appealing that the prolonged inhibition of transcriptional activity leads to the formation of morphologically similar compact domains in IGC/nuclear speckles as neuronal stimulation. However, their physiological resemblance and function have not been studied yet.

Contrary to transcriptional inhibition, blocking of splicing in neurons increased the number of compact domains, even in the absence of stimulation. Several authors have noticed structural changes to nuclear speckles as a result of splicing inhibition. O’Keefe et al. showed that microinjection of oligonucleotides blocking the action of splicing factors U1 and U6 snRNP resulted in larger and more rounded IGCs/nuclear speckles (O’Keefe et al., 1994). However, the electron micrograph presented by the authors did not show any distinguishable aggregates in IGCs. Kaida et al. observed a similar effect of splicing inhibition by spliceostatin A (Kaida et al., 2007). The enlargement of nuclear speckles was accompanied by increased levels of poly(A)+ mRNAs in the nuclei. The authors concluded that the observed structural changes of the nuclear speckles are a result of the accumulation of transcripts incapable of correct splicing. More precisely, Carvalho et al. showed retention of the unspliced β-globin and β-actin pre-mRNAs in the enlarged nuclear speckles as a result of the splicing inhibition with spliceostatin A, meayamycin, or pladienolide B (Carvalho et al., 2017). Unfortunately, none of the aforementioned articles presented ultrastructural data.

### 4.2. Changes in protein composition of nuclear speckles upon neuronal activation

Previous studies of the nuclear speckle proteome were performed on mouse hepatocytes (Saitoh et al., 2004) and the U2OS cell line (Dopie et al., 2020). Similarly to those studies, our analysis has shown that the neuronal speckle proteome is dominated by factors involved in mRNA processing, mainly splicing factors. Both comparative (qualitative) and quantitative analysis of IGC/nuclear speckles composition showed that neuronal stimulation increases levels of many proteins from the hnRNP family. This may be related to the ultrastructural changes of IGC/nuclear speckles observed upon neuronal stimulation. The hnRNPs are responsible for almost every stage of mRNA processing in the nucleus regulating mRNA assembly, mRNA packing into hnRNP particles, their stabilization, and export to the cytoplasm (Jean-Philippe et al., 2013; Liu and Shi, 2021). As already mentioned, the stimulation of neurons induces massive activation of transcription of hundreds of genes involved in neuronal plasticity (Fernandez-Albert et al., 2019; Grabowska et al., 2022). It seems possible that in response to activation, neuronal cells increase levels of proteins residing in nuclear speckles, involved in processing of newly produced mRNAs. Moreover, the accumulating splicing factors and the RNA products of the activated genes may also induce structural changes in IGC/nuclear speckles. We showed that neuronal stimulation induced the assembly of very large nuclear speckles in the range of 13.5 µm^3^ - 18.5 µm^3^ and increased the volume of compact domains. Furthermore, inhibition of splicing and in consequence accumulation of the unspliced mRNAs, amplified the occurrence of the compact domains and, in line with others authors, resulted in larger and more rounded IGCs/nuclear speckles (Carvalho et al., 2017; Kaida et al., 2007; O’Keefe et al., 1994). In addition, we observed increased levels of SFPQ and SF3A2 proteins localized in compact domains of stimulated neurons. Both proteins are essential components of the spliceosome (Gozani et al., 1994; Nesic and Kramer, 2001; Will and Luhrmann, 2011). Therefore, their increased localization in compact domains may indicate activity-induced accumulation of the spliceosomes together with processed pre-mRNAs. Unfortunately, our attempts to verify the presence of the hnRNPs directly at the ultrastructural level were unsuccessful as the immunogold detection of hnRNPs showed no or very poor labelling, in the IGC/nuclear speckles, both in control and stimulated neurons (data not shown). The high concentrations of hnRNP proteins in nuclear speckles may cause their partial oligomerization and/or aggregation (Deshaies et al., 2018; Kim et al., 2013) and consequently may lead to the so-called steric hindrance preventing attachment of the antibodies, especially those conjugated with relatively large colloidal gold particles (Amiry-Moghaddam and Ottersen, 2013).

While transcription factors in studies of Saitoh and Dopie (Dopie et al., 2020; Saitoh et al., 2004) represented only about 3% and 11%, respectively, of all proteins detected in the nuclear speckles from non-neuronal cells, we showed, that it was the second largest functional group constituting 36% of all proteins found in neuronal speckles. In addition, the levels of 13 proteins involved in transcription increased in nuclear speckles, upon neuronal stimulation. Further, our results demonstrated the presence of DNA and the activated forms of RNAPII. The levels of the phosphorylated forms of RNAPII were also increased in nuclear speckles upon neuronal activation, a known trigger of a massive transcriptional boost. We also noticed a disappearance of compact domains upon inhibition of transcription. All these results might suggest an involvement of neuronal nuclear speckles, and possibly of compact domains, in regulation of genes transcription induced by neuronal activity. The presence of RNAPII in nuclear speckles of MDCK cells was first reported by Bregman et al. (Bregman et al., 1995). Another study by Xie et al. showed that nuclear speckles of HeLa cells contain both forms of activated RNAPII but their levels were not higher than in the nucleoplasm. However, the authors suggested that the presence of a stably bound, small pool of RNAPII phosphorylated on Ser2 in nuclear speckles was not associated with transcriptional activity, but rather with the formation of the spliceosome and splicing initiation (Xie et al., 2006), for which RNAPII is mandatory (Hirose et al., 1999; Zeng and Berget, 2000). On the other hand, an association of actively transcribed genes with the nuclear speckles was shown in many models (Brown et al., 2006; Chen et al., 2018; Ding and Elowitz, 2019; Guo et al., 2019; Kim et al., 2020; Zhang et al., 2021), where levels and location of transcription and splicing positively correlated with proximity to nuclear speckles. In a recent publication, Alexander et al. showed that the association of genes with nuclear speckles can be driven by the p53 transcription factor, and such association elevated the production of the nascent RNAs and boosted gene expression (Alexander et al., 2021). The p53-dependent genes, which were not localized in the nuclear speckles vicinity had lower levels of expression than p53-dependent genes which were associated with those structures. Moreover, the knockdown of SON and SRSF1, components involved in nuclear speckles organization, reduced expression of p21, dependent on p53. This shows that the structural integrity and composition of nuclear speckles may also influence gene expression. Some authors suggested that nuclear speckles are formed at the sites of active transcription (Brown et al., 2008; Guo et al., 2019; Hu et al., 2010). Although the total number of nuclear speckles was not significantly changed upon the kainate-induced neuronal stimulation *in vivo*, the smallest nuclear speckles (0.5-1.5 µm^3^) were more abundant compared to the unstimulated neurons. Moreover, in the *in vitro* model, activation through chemically-evoked LTP significantly increased the number of nuclear speckles. Most of the nuclear speckles resided in enlarged interchromatin space, that appears as a result of the activity-dependent chromatin condensation (Grabowska et al., 2022). It is therefore possible that the stimulation of neurons and the resulting expression boost of neuronal genes induce formation of new nuclear speckles at the sites of active transcription. Interestingly, the localization of DNA within nuclear speckles was more central in activated neurons, compared to the control cells, where it was found close to the speckle surface.

## 5. Conclusion

Taking into account the above results, we propose that the compact domains induced by neuronal stimulation are the ultrastructural manifestation of the accumulation of newly transcribed mRNAs together with proteins present in IGC/nuclear speckles. The load of transcripts involved in neuronal plasticity rapidly increases upon the stimulation and undergoes a gradual accumulation within nuclear speckles for further processing and later export (Fig. 12).

**Figure 12.**
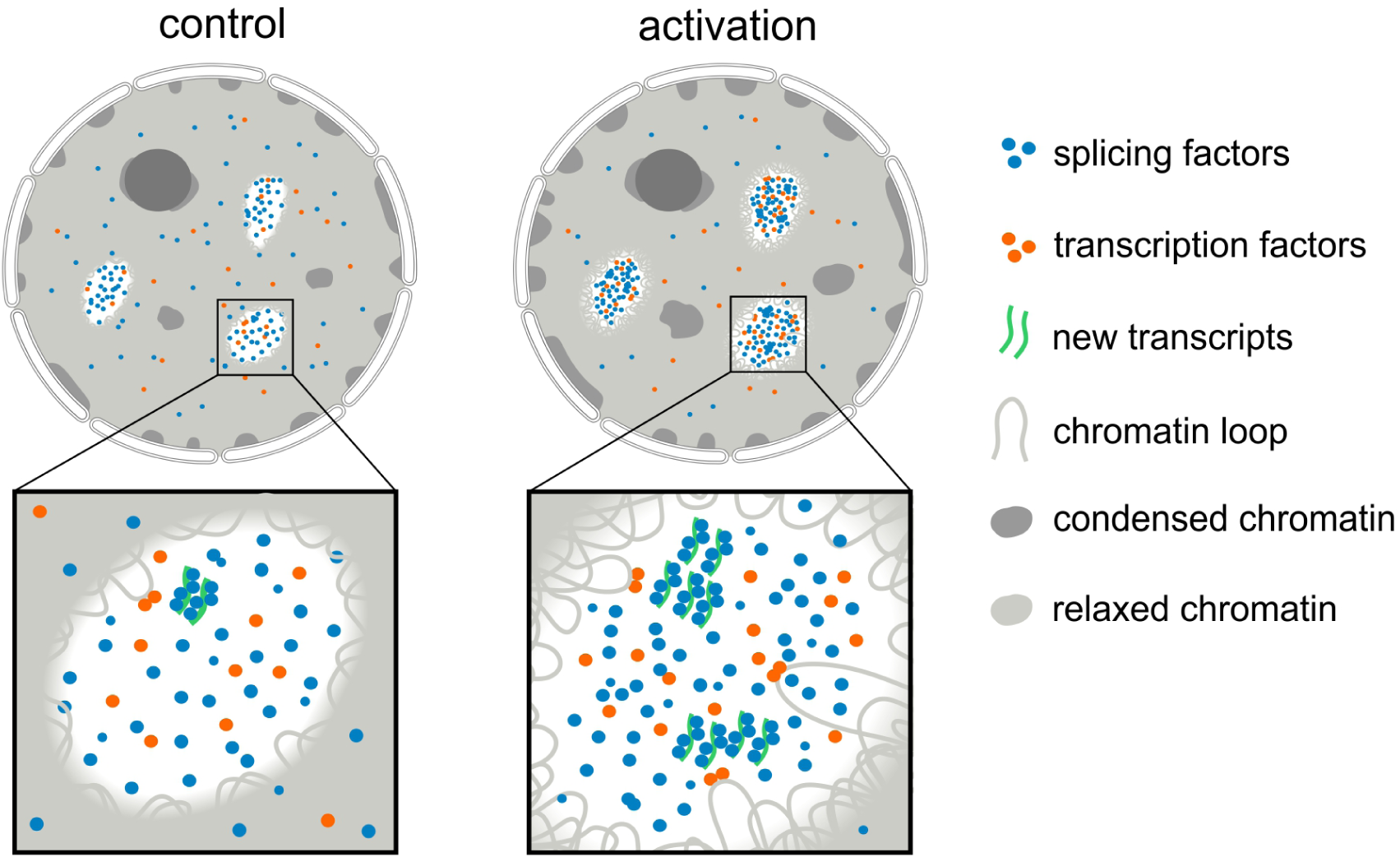
Neuronal nuclear speckles upon activation. Neuronal activation induces the transcription of genes involved e.g. in neuronal plasticity. According to our model, newly created transcripts undergo accumulation in the IGCs/nuclear speckles for more efficient processing and export. The manifestation of this process is the assembly of the compact domains observed in the IGCs/nuclear speckles at the ultrastructural level. It is accompanied by the accumulation of DNA and specific proteins in the ICGs/nuclear speckles.

## Supporting information

Supplementary MassSpec Data

## 6. CRediT authorship contribution statement

### Andrzej Antoni Szczepankiewicz

Funding acquisition, Conceptualization, Resources, Methodology, Investigation, Visualization, Formal analysis, Writing – original draft, Writing – review & editing.

### Kamil Parobczak

Investigation, Formal analysis, Writing – review & editing.

### Monika Zaręba-Kozioł

Investigation, Formal analysis.

### Błażej Ruszczycki

Software, Formal analysis, Writing – review & editing.

### Monika Bijata

Investigation, Resources.

### Paweł Trzaskoma

Investigation, Writing – review & editing.

### Grzegorz Hajnowski

Investigation.

### Dagmara Holm-Kaczmarek

Investigation.

### Jakub Włodarczyk

Funding acquisition, Supervision.

### Grzegorz Marek Wilczyński

Conceptualization, Funding acquisition, Methodology, Supervision, Project administration.

### Maria Jolanta Rędowicz

Writing – review & editing, Supervision.

### Adriana Magalska

Writing – original draft, Writing – review & editing, Supervision.

## 7. Acknowledgments

This work was supported primarily by the Polish National Science Centre grant UMO-2014/15/N/NZ3/04468. Monika Bijata, Monika Zaręba-Kozioł and Jakub Włodarczyk were supported by the Polish National Science Centre grants UMO-2021/41/B/NZ4/02603 and UMO-2019/35/D/NZ4/02042. We are grateful to Iwona Czaban, Hubert Doleżyczek, Małgorzata Hall, Elżbieta Januszewicz, Katarzyna Pels and Agnieszka Walczak for their help in kainic acid experiments for microscopic analysis. We thank Matylda Macias and Magdalena Błażejczyk for their help in kainic acid experiments for mass spectrometry analysis. We would like to acknowledge also Tomasz Górkiewicz, Robert Kuba Filipkowski and Monika Malinowska for helping in experiments that were ultimately not included in this work. We thank David Spector (Cold Spring Harbor Laboratory, NY, USA) for sharing the anti-phospho-SR (3C5) antibody.

## 8. References

Alexander, K.A., A. Cote, S.C. Nguyen, L. Zhang, O. Gholamalamdari, P. Agudelo-Garcia, E. Lin-Shiao, K.M.A. Tanim, J. Lim, N. Biddle, M.C. Dunagin, C.R. Good, M.R. Mendoza, S.C. Little, A. Belmont, E.F. Joyce, A. Raj, and S.L. Berger. 2021. p53 mediates target gene association with nuclear speckles for amplified RNA expression. Mol Cell.

Amiry-Moghaddam, M., and O.P. Ottersen. 2013. Immunogold cytochemistry in neuroscience. Nat Neurosci. 16:798–804.

Barutcu, A.R., M. Wu, U. Braunschweig, B.J.A. Dyakov, Z. Luo, K.M. Turner, T. Durbic, Z.Y. Lin, R.J. Weatheritt, P.G. Maass, A.C. Gingras, and B.J. Blencowe. 2022. Systematic mapping of nuclear domain-associated transcripts reveals speckles and lamina as hubs of functionally distinct retained introns. Mol Cell. 82:1035–1052 e1039.

Beck, J.S. 1961. Variations in the Morphological Patterns of “Autoimmune” Nuclear Fluorescence. The Lancet. 277:1203–1205.

Ben-Ari, Y. 1985. Limbic seizure and brain damage produced by kainic acid: mechanisms and relevance to human temporal lobe epilepsy. Neuroscience. 14:375–403.

Bhat, P., A. Chow, B. Emert, O. Ettlin, S.A. Quinodoz, Y. Takei, W. Huang, M.R. Blanco, and M. Guttman. 2023. 3D genome organization around nuclear speckles drives mRNA splicing efficiency. bioRxiv.

Bregman, D.B., L. Du, S. van der Zee, and S.L. Warren. 1995. Transcription-dependent redistribution of the large subunit of RNA polymerase II to discrete nuclear domains. J. Cell Biol. 129:287–298.

Brown, J.M., J. Green, R.P.D. Neves, H.A.C. Wallace, A.J.H. Smith, J. Hughes, N. Gray, S. Taylor, W.G. Wood, D.R. Higgs, F.J. Iborra, and V.J. Buckle. 2008. Association between active genes occurs at nuclear speckles and is modulated by chromatin environment. Journal of Cell Biology. 182:1083–1097.

Brown, J.M., J. Leach, J.E. Reittie, A. Atzberger, J. Lee-Prudhoe, W.G. Wood, D.R. Higgs, F.J. Iborra, and V.J. Buckle. 2006. Coregulated human globin genes are frequently in spatial proximity when active. J Cell Biol. 172:177–187.

Cajal, S.R. 1910. El núcleo de las células piramidales del cerebro humano y de algunos mamíferos. Trab Lab Invest Biol. 8:35.

Carvalho, T., S. Martins, J. Rino, S. Marinho, and M. Carmo-Fonseca. 2017. Pharmacological inhibition of the spliceosome subunit SF3b triggers exon junction complex-independent nonsense-mediated decay. J Cell Sci. 130:1519–1531.

Chen, Y., Y. Zhang, Y. Wang, L. Zhang, E.K. Brinkman, S.A. Adam, R. Goldman, B. van Steensel, J. Ma, and A.S. Belmont. 2018. Mapping 3D genome organization relative to nuclear compartments using TSA-Seq as a cytological ruler. J Cell Biol. 217:4025–4048.

Deshaies, J.E., L. Shkreta, A.J. Moszczynski, H. Sidibe, S. Semmler, A. Fouillen, E.R. Bennett, U. Bekenstein, L. Destroismaisons, J. Toutant, Q. Delmotte, K. Volkening, S. Stabile, A. Aulas, Y. Khalfallah, H. Soreq, A. Nanci, M.J. Strong, B. Chabot, and C. Vande Velde. 2018. TDP-43 regulates the alternative splicing of hnRNP A1 to yield an aggregation-prone variant in amyotrophic lateral sclerosis. Brain. 141:1320–1333.

Dias, A.P., K. Dufu, H. Lei, and R. Reed. 2010. A role for TREX components in the release of spliced mRNA from nuclear speckle domains. Nat Commun. 1:97.

Ding, F., and M.B. Elowitz. 2019. Constitutive splicing and economies of scale in gene expression. Nat Struct Mol Biol. 26:424–432.

Dopie, J., M.J. Sweredoski, A. Moradian, and A.S. Belmont. 2020. Tyramide signal amplification mass spectrometry (TSA-MS) ratio identifies nuclear speckle proteins. The Journal of cell biology. 219.

Dragunow, M., and H.A. Robertson. 1987. Generalized seizures induce c-fos protein(s) in mammalian neurons. Neurosci Lett. 82:157–161.

Elias, H., and D.M. Hyde. 1980. An elementary introduction to stereology (quantitative microscopy). Am J Anat. 159:412–446.

Fedorko, M.E., and J.G. Hirsch. 1969. Nucleolar fragmentation in L cells exposed to quinacrine in vitro. Cancer Res. 29:918–924.

Fernandez-Albert, J., M. Lipinski, M.T. Lopez-Cascales, M.J. Rowley, A.M. Martin-Gonzalez, B. Del Blanco, V.G. Corces, and A. Barco. 2019. Immediate and deferred epigenomic signatures of in vivo neuronal activation in mouse hippocampus. Nat Neurosci. 22:1718–1730.

Fox, A.H., and A.I. Lamond. 2010. Paraspeckles. Cold Spring Harb Perspect Biol. 2:a000687.

Galganski, L., M.O. Urbanek, and W.J. Krzyzosiak. 2017. Nuclear speckles: molecular organization, biological function and role in disease. Nucleic Acids Res. 45:10350–10368.

Gavrieli, Y., Y. Sherman, and S.A. Ben-Sasson. 1992. Identification of programmed cell death in situ via specific labeling of nuclear DNA fragmentation. The Journal of cell biology. 119:493–501.

Girard, C., C.L. Will, J. Peng, E.M. Makarov, B. Kastner, I. Lemm, H. Urlaub, K. Hartmuth, and R. Luhrmann. 2012. Post-transcriptional spliceosomes are retained in nuclear speckles until splicing completion. Nat Commun. 3:994.

Gozani, O., J.G. Patton, and R. Reed. 1994. A novel set of spliceosome-associated proteins and the essential splicing factor PSF bind stably to pre-mRNA prior to catalytic step II of the splicing reaction. The EMBO journal. 13:3356–3367.

Grabowska, A., H. Sas-Nowosielska, B. Wojtas, D. Holm-Kaczmarek, E. Januszewicz, Y. Yushkevich, I. Czaban, P. Trzaskoma, K. Krawczyk, B. Gielniewski, A. Martin-Gonzalez, R.K. Filipkowski, K.H. Olszynski, T. Bernas, A.A. Szczepankiewicz, M.A. Sliwinska, T. Kanhema, C.R. Bramham, G. Bokota, D. Plewczynski, G.M. Wilczynski, and A. Magalska. 2022. Activation-induced chromatin reorganization in neurons depends on HDAC1 activity. Cell Rep. 38:110352.

Griffiths, G. 1993. Fine Structure Immunocytochemistry. Springer-Verlag Berlin Heidelberg, Heidelberg.

Guo, Y.E., J.C. Manteiga, J.E. Henninger, B.R. Sabari, A. Dall’Agnese, N.M. Hannett, J.H. Spille, L.K. Afeyan, A.V. Zamudio, K. Shrinivas, B.J. Abraham, A. Boija, T.M. Decker, J.K. Rimel, C.B. Fant, T.I. Lee, Cisse, II, P.A. Sharp, D.J. Taatjes, and R.A. Young. 2019. Pol II phosphorylation regulates a switch between transcriptional and splicing condensates. Nature. 572:543–548.

Hall, L.L., K.P. Smith, M. Byron, and J.B. Lawrence. 2006. Molecular anatomy of a speckle. *Anatomical Record - Part A Discoveries in Molecular*, Cellular, and Evolutionary Biology. 288:664–675.

Hall, M.H., A. Magalska, M. Malinowska, B. Ruszczycki, I. Czaban, S. Patel, M. Ambrozek-Latecka, E. Zolocinska, H. Broszkiewicz, K. Parobczak, R.R. Nair, M. Rylski, R. Pawlak, C.R. Bramham, and G.M. Wilczynski. 2016. Localization and regulation of PML bodies in the adult mouse brain. Brain Struct Funct. 221:2511–2525.

Hattinger, C.M., A.G. Jochemsen, H.J. Tanke, and R.W. Dirks. 2002. Induction of p21 mRNA synthesis after short-wavelength UV light visualized in individual cells by RNA FISH. J Histochem Cytochem. 50:81–89.

Hellier, J.L., P.R. Patrylo, P.S. Buckmaster, and F.E. Dudek. 1998. Recurrent spontaneous motor seizures after repeated low-dose systemic treatment with kainate: assessment of a rat model of temporal lobe epilepsy. Epilepsy Res. 31:73–84.

Hirose, Y., R. Tacke, and J.L. Manley. 1999. Phosphorylated RNA polymerase II stimulates pre-mRNA splicing. Genes Dev. 13:1234–1239.

Hu, Y., M. Plutz, and A.S. Belmont. 2010. Hsp70 gene association with nuclear speckles is Hsp70 promoter specific. The Journal of cell biology. 191:711–719.

Huang, S., A. Mayeda, A.R. Krainer, and D.L. Spector. 1997. RCC1 and nuclear organization. Molecular biology of the cell. 8:1143–1157.

Huang, Y., J.J. Doherty, and R. Dingledine. 2002. Altered histone acetylation at glutamate receptor 2 and brain-derived neurotrophic factor genes is an early event triggered by status epilepticus. J Neurosci. 22:8422–8428.

Ikenari, T., H. Kurata, T. Satoh, Y. Hata, and T. Mori. 2020. Evaluation of Fluoro-Jade C Staining: Specificity and Application to Damaged Immature Neuronal Cells in the Normal and Injured Mouse Brain. Neuroscience. 425:146–156.

Jean-Philippe, J., S. Paz, and M. Caputi. 2013. hnRNP A1: the Swiss army knife of gene expression. Int J Mol Sci. 14:18999–19024.

Johnson, C. 2000. Tracking COL1A1 RNA in osteogenesis imperfecta. Splice-defective transcripts initiate transport from the gene but are retained within the SC35 domain. J. Cell Biol. 150:417–432.

Jolly, C., C. Vourc’h, M. Robert-Nicoud, and R.I. Morimoto. 1999. Intron-independent association of splicing factors with active genes. J Cell Biol. 145:1133–1143.

Jones, K.J., and A. LaVelle. 1986. Ultrastructural changes in the nucleoplasm of hamster facial neurons during a postnatal maturation period. Brain research. 377:119–126.

Kaida, D., H. Motoyoshi, E. Tashiro, T. Nojima, M. Hagiwara, K. Ishigami, H. Watanabe, T. Kitahara, T. Yoshida, H. Nakajima, T. Tani, S. Horinouchi, and M. Yoshida. 2007. Spliceostatin A targets SF3b and inhibits both splicing and nuclear retention of pre-mRNA. Nat Chem Biol. 3:576–583.

Kim, H.J., N.C. Kim, Y.D. Wang, E.A. Scarborough, J. Moore, Z. Diaz, K.S. MacLea, B. Freibaum, S. Li, A. Molliex, A.P. Kanagaraj, R. Carter, K.B. Boylan, A.M. Wojtas, R. Rademakers, J.L. Pinkus, S.A. Greenberg, J.Q. Trojanowski, B.J. Traynor, B.N. Smith, S. Topp, A.S. Gkazi, J. Miller, C.E. Shaw, M. Kottlors, J. Kirschner, A. Pestronk, Y.R. Li, A.F. Ford, A.D. Gitler, M. Benatar, O.D. King, V.E. Kimonis, E.D. Ross, C.C. Weihl, J. Shorter, and J.P. Taylor. 2013. Mutations in prion-like domains in hnRNPA2B1 and hnRNPA1 cause multisystem proteinopathy and ALS. Nature. 495:467–473.

Kim, J., K.Y. Han, N. Khanna, T. Ha, and A.S. Belmont. 2019. Nuclear speckle fusion via long-range directional motion regulates speckle morphology after transcriptional inhibition. J Cell Sci. 132.

Kim, J., N.C. Venkata, G.A. Hernandez Gonzalez, N. Khanna, and A.S. Belmont. 2020. Gene expression amplification by nuclear speckle association. The Journal of cell biology. 219.

Krzyzowska-Gruca, S., A. Zborek, and S. Gruca. 1983. Distribution of interchromatin granules in nuclear matrices obtained from nuclei exhibiting different degree of chromatin condensation. Cell Tissue Res. 231:427–437.

Laemmli, U.K. 1970. Cleavage of structural proteins during the assembly of the head of bacteriophage T4. Nature. 227:680–685.

Lafarga, M., M.T. Berciano, L.M. Garcia-Segura, M.A. Andres, and M. Carmo-Fonseca. 1998. Acute osmotic/stress stimuli induce a transient decrease of transcriptional activity in the neurosecretory neurons of supraoptic nuclei. Journal of neurocytology. 27:205–217.

Lafarga, M., O. Tapia, A.M. Romero, and M.T. Berciano. 2017. Cajal bodies in neurons. RNA Biol. 14:712–725.

Letunic, I., and P. Bork. 2018. 20 years of the SMART protein domain annotation resource. Nucleic acids research. 46:D493–D496.

Liu, Y., and S.L. Shi. 2021. The roles of hnRNP A2/B1 in RNA biology and disease. Wiley interdisciplinary reviews. RNA. 12:e1612.

Locke, M., and P. Huie. 1977. Bismuth staining for light and electron microscopy. Tissue Cell. 9:347–371.

Malinowska, A., M. Kistowski, M. Bakun, T. Rubel, M. Tkaczyk, J. Mierzejewska, and M. Dadlez. 2012. Diffprot - software for non-parametric statistical analysis of differential proteomics data. J Proteomics. 75:4062–4073.

Marinozzi, V., and L. Fiume. 1971. Effects of -amanitin on mouse and rat liver cell nuclei. Experimental cell research. 67:311–322.

Matevossian, A., and S. Akbarian. 2008. Neuronal nuclei isolation from human postmortem brain tissue. J Vis Exp.

Mathiisen, T.M., E.A. Nagelhus, B. Jouleh, R. Torp, D.S. Frydenlund, M.-N. Mylonakou, M. Amiry-Moghaddam, L. Covolan, J.K. Utvik, B. Riber, K.M. Gujord, J. Knutsen, Ø. Skare, P. Laake, S. Davanger, F.-M. Haug, E. Rinvik, and O.P. Ottersen. 2006. Postembedding Immunogold Cytochemistry of Membrane Molecules and Amino Acid Transmitters in the Central Nervous System. In Neuroanatomical Tract-Tracing 3: Molecules, Neurons, and Systems. L. Zaborszky, F.G. Wouterlood, and J.L. Lanciego, editors. Springer US, Boston, MA. 72–108.

Melcak, I., S. Cermanova, K. Jirsova, K. Koberna, J. Malinsky, and I. Raska. 2000. Nuclear pre-mRNA compartmentalization: trafficking of released transcripts to splicing factor reservoirs. Mol Biol Cell. 11:497–510.

Mintz, P.J., S.D. Patterson, A.F. Neuwald, C.S. Spahr, and D.L. Spector. 1999. Purification and biochemical characterization of interchromatin granule clusters. EMBO J. 18:4308–4320.

Miyai, K., and J.W. Steiner. 1965. Fine structure of interphase liver cell nuclei in subacute ethionine intoxication. Exp Mol Pathol. 4:525–566.

Moen Jr, P.T., C.V. Johnson, M. Byron, L.S. Shopland, I.L. De La Serna, A.N. Imbalzano, and J.B. Lawrence. 2004. Repositioning of Muscle-specific Genes Relative to the Periphery of SC-35 Domains during Skeletal Myogenesis. Molecular Biology of the Cell. 15:197–206.

Morgan, J.I., D.R. Cohen, J.L. Hempstead, and T. Curran. 1987. Mapping patterns of c-fos expression in the central nervous system after seizure. Science. 237:192–197.

Morris, G.E. 2008. The Cajal body. Biochim Biophys Acta. 1783:2108–2115.

Naum-Ongania, G., V.M. Diaz, F. Blasi, and R. Rivera-Pomar. 2013. Nuclear actin polymerization from faster growing ends in the initial activation of Hox gene transcription are nuclear speckles involved? Transcription. 4:260–272.

Navascues, J., I. Casafont, N.T. Villagra, M. Lafarga, and M.T. Berciano. 2004. Reorganization of nuclear compartments of type A neurons of trigeminal ganglia in response to inflammatory injury of peripheral nerve endings. J Neurocytol. 33:393–405.

Nesic, D., and A. Kramer. 2001. Domains in human splicing factors SF3a60 and SF3a66 required for binding to SF3a120, assembly of the 17S U2 snRNP, and prespliceosome formation. Molecular and cellular biology. 21:6406–6417.

Niedojadlo, J., Z. Mikulski, K. Delenko, A. Szmidt-Jaworska, D.J. Smolinski, and A.L. Epstein. 2012. The perichromatin region of the plant cell nucleus is the area with the strongest co-localisation of snRNA and SR proteins. Planta. 236:715–726.

Nielsen, J.A., L.D. Hudson, and R.C. Armstrong. 2002. Nuclear organization in differentiating oligodendrocytes. J Cell Sci. 115:4071–4079.

O’Keefe, R.T., A. Mayeda, C.L. Sadowski, A.R. Krainer, and D.L. Spector. 1994. Disruption of pre-mRNA splicing in vivo results in reorganization of splicing factors. In Journal of Cell Biology. Vol. 124. 249–260.

Obay, B.D., E. Tasdemir, C. Tumer, H.M. Bilgin, and A. Sermet. 2007. Antiepileptic effects of ghrelin on pentylenetetrazole-induced seizures in rats. Peptides. 28:1214–1219.

Otmakhov, N., L. Khibnik, N. Otmakhova, S. Carpenter, S. Riahi, B. Asrican, and J. Lisman. 2004. Forskolin-induced LTP in the CA1 hippocampal region is NMDA receptor dependent. J Neurophysiol. 91:1955–1962.

Pena, E., M.T. Berciano, R. Fernandez, J.L. Ojeda, and M. Lafarga. 2001. Neuronal body size correlates with the number of nucleoli and Cajal bodies, and with the organization of the splicing machinery in rat trigeminal ganglion neurons. J Comp Neurol. 430:250–263.

Pranchevicius, M.C., M.M. Baqui, H.C. Ishikawa-Ankerhold, E.V. Lourenco, R.M. Leao, S.R. Banzi, C.T. dos Santos, M.C. Roque-Barreira, E.M. Espreafico, and R.E. Larson. 2008. Myosin Va phosphorylated on Ser1650 is found in nuclear speckles and redistributes to nucleoli upon inhibition of transcription. Cell Motil Cytoskeleton. 65:441–456.

Quinodoz, S.A., N. Ollikainen, B. Tabak, A. Palla, J.M. Schmidt, E. Detmar, M.M. Lai, A.A. Shishkin, P. Bhat, Y. Takei, V. Trinh, E. Aznauryan, P. Russell, C. Cheng, M. Jovanovic, A. Chow, L. Cai, P. McDonel, M. Garber, and M. Guttman. 2018. Higher-Order Inter-chromosomal Hubs Shape 3D Genome Organization in the Nucleus. Cell. 174:744–757 e724.

Racine, R.J. 1972. Modification of seizure activity by electrical stimulation. II. Motor seizure. Electroencephalogr Clin Neurophysiol. 32:281–294.

Reynolds, R.C., P.O. Montgomery, and B. Hughes. 1964. Nucleolar “Caps” Produced by Actinomycin D. Cancer Res. 24:1269–1277.

Ruszczycki, B., K.K. Pels, A. Walczak, K. Zamlynska, M. Such, A.A. Szczepankiewicz, M.H. Hall, A. Magalska, M. Magnowska, A. Wolny, G. Bokota, S. Basu, A. Pal, D. Plewczynski, and G.M. Wilczynski. 2019. Three-Dimensional Segmentation and Reconstruction of Neuronal Nuclei in Confocal Microscopic Images. Front Neuroanat. 13:81.

Saitoh, N., C.S. Spahr, S.D. Patterson, P. Bubulya, A.F. Neuwald, and D.L. Spector. 2004. Proteomic analysis of interchromatin granule clusters. Molecular Biology of the Cell. 15:3876–3890.

Sawicka, A., and C. Seiser. 2012. Histone H3 phosphorylation - a versatile chromatin modification for different occasions. Biochimie. 94:2193–2201.

Schindelin, J., I. Arganda-Carreras, E. Frise, V. Kaynig, M. Longair, T. Pietzsch, S. Preibisch, C. Rueden, S. Saalfeld, B. Schmid, J.Y. Tinevez, D.J. White, V. Hartenstein, K. Eliceiri, P. Tomancak, and A. Cardona. 2012. Fiji: an open-source platform for biological-image analysis. Nat Methods. 9:676–682.

Schoefl, G.I. 1964. The Effect of Actinomycin D on the Fine Structure of the Nucleolus. J Ultrastruct Res. 10:224–243.

Shopland, L.S., C.V. Johnson, and J.B. Lawrence. 2002. Evidence that all SC-35 domains contain mRNAs and that transcripts can be structurally constrained within these domains. J Struct Biol. 140:131–139.

Skupien-Jaroszek, A., A. Walczak, I. Czaban, K.K. Pels, A.A. Szczepankiewicz, K. Krawczyk, B. Ruszczycki, G.M. Wilczynski, J. Dzwonek, and A. Magalska. 2021. The interplay of seizures-induced axonal sprouting and transcription-dependent Bdnf repositioning in the model of temporal lobe epilepsy. PLoS One. 16:e0239111.

Smith, K.P., P.T. Moen, K.L. Wydner, J.R. Coleman, and J.B. Lawrence. 1999. Processing of endogenous pre-mRNAs in association with SC-35 domains is gene specific. J. Cell Biol. 144:617–629.

Spector, D.L., and A.I. Lamond. 2011. Nuclear speckles. Cold Spring Harb Perspect Biol. 3.

Spector, D.L., R.T. O’Keefe, and L.F. Jimenez-Garcia. 1993. Dynamics of transcription and pre-mRNA splicing within the mammalian cell nucleus. Cold Spring Harb Symp Quant Biol. 58:799–805.

Sperk, G. 1994. Kainic acid seizures in the rat. Prog Neurobiol. 42:1–32.

Stoppini, L., P.A. Buchs, and D. Muller. 1991. A simple method for organotypic cultures of nervous tissue. J Neurosci Methods. 37:173–182.

Svoboda, D., and J. Higginson. 1968. A comparison of ultrastructural changes in rat liver due to chemical carcinogens. Cancer Res. 28:1703–1733.

Swift, H. 1959. Studies on nuclear fine structure. Brookhaven Symp Biol. 12:134–152.

Szczerbal, I., and J.M. Bridger. 2010. Association of adipogenic genes with SC-35 domains during porcine adipogenesis. Chromosome Res. 18:887–895.

Szepesi, Z., E. Hosy, B. Ruszczycki, M. Bijata, M. Pyskaty, A. Bikbaev, M. Heine, D. Choquet, L. Kaczmarek, and J. Wlodarczyk. 2014. Synaptically released matrix metalloproteinase activity in control of structural plasticity and the cell surface distribution of GluA1-AMPA receptors. PLoS One. 9:e98274.

Taniura, H., J.C. Sng, and Y. Yoneda. 2006. Histone modifications in status epilepticus induced by kainate. Histology and histopathology. 21:785–791.

Tao-Cheng, J.H. 2018. Stimulation-induced structural changes at the nucleus, endoplasmic reticulum and mitochondria of hippocampal neurons. Mol Brain. 11:44.

Thiry, M. 1992. Highly sensitive immunodetection of DNA on sections with exogenous terminal deoxynucleotidyl transferase and non-isotopic nucleotide analogues. The journal of histochemistry and cytochemistry: official journal of the Histochemistry Society. 40:411–419.

Tripathi, V., D.Y. Song, X. Zong, S.P. Shevtsov, S. Hearn, X.D. Fu, M. Dundr, and K.V. Prasanth. 2012. SRSF1 regulates the assembly of pre-mRNA processing factors in nuclear speckles. Molecular biology of the cell. 23:3694–3706.

Turner, B.M., and S. Davies. 1986. A monoclonal antibody recognizes a phosphorylated epitope shared by proteins of the cell nucleus and the erythrocyte membrane skeleton. FEBS letters. 197:41–44.

van der Lee, R., M. Buljan, B. Lang, R.J. Weatheritt, G.W. Daughdrill, A.K. Dunker, M. Fuxreiter, J. Gough, J. Gsponer, D.T. Jones, P.M. Kim, R.W. Kriwacki, C.J. Oldfield, R.V. Pappu, P. Tompa, V.N. Uversky, P.E. Wright, and M.M. Babu. 2014. Classification of intrinsically disordered regions and proteins. Chem Rev. 114:6589–6631.

Walczak, A., A.A. Szczepankiewicz, B. Ruszczycki, A. Magalska, K. Zamlynska, J. Dzwonek, E. Wilczek, K. Zybura-Broda, M. Rylski, M. Malinowska, M. Dabrowski, T. Szczepinska, K. Pawlowski, M. Pyskaty, J. Wlodarczyk, I. Szczerbal, M. Switonski, M. Cremer, and G.M. Wilczynski. 2013. Novel higher-order epigenetic regulation of the Bdnf gene upon seizures. J Neurosci. 33:2507–2511.

Wang, K., L. Wang, J. Wang, S. Chen, M. Shi, and H. Cheng. 2018. Intronless mRNAs transit through nuclear speckles to gain export competence. J Cell Biol. 217:3912–3929.

Will, C.L., and R. Luhrmann. 2011. Spliceosome structure and function. Cold Spring Harbor perspectives in biology. 3.

Xie, S.Q., S. Martin, P.V. Guillot, D.L. Bentley, and A. Pombo. 2006. Splicing speckles are not reservoirs of RNA polymerase II, but contain an inactive form, phosphorylated on serine2 residues of the C-terminal domain. Mol Biol Cell. 17:1723–1733.

Xing, Y., C.V. Johnson, P.T. Moen, J.A. McNeil, and J.B. Lawrence. 1995. Nonrandom gene organization: structural arrangements of specific pre-mRNA transcription and splicing with SC35 domains. J. Cell Biol. 131:1635–1647.

Yarosh, C.A., J.R. Iacona, C.S. Lutz, and K.W. Lynch. 2015. PSF: nuclear busy-body or nuclear facilitator? Wiley interdisciplinary reviews. RNA. 6:351–367.

Yokoi, A., Y. Kotake, K. Takahashi, T. Kadowaki, Y. Matsumoto, Y. Minoshima, N.H. Sugi, K. Sagane, M. Hamaguchi, M. Iwata, and Y. Mizui. 2011. Biological validation that SF3b is a target of the antitumor macrolide pladienolide. Febs J. 278:4870–4880.

Zagulska-Szymczak, S., R.K. Filipkowski, and L. Kaczmarek. 2001. Kainate-induced genes in the hippocampus: lessons from expression patterns. Neurochem Int. 38:485–501.

Zareba-Koziol, M., A. Bartkowiak-Kaczmarek, I. Figiel, A. Krzystyniak, T. Wojtowicz, M. Bijata, and J. Wlodarczyk. 2019. Stress-induced Changes in the S-palmitoylation and S-nitrosylation of Synaptic Proteins. Mol Cell Proteomics. 18:1916–1938.

Zeng, C., and S.M. Berget. 2000. Participation of the C-terminal domain of RNA polymerase II in exon definition during pre-mRNA splicing. Mol Cell Biol. 20:8290–8301.

Zhang, L., Y. Zhang, Y. Chen, O. Gholamalamdari, Y. Wang, J. Ma, and A.S. Belmont. 2021. TSA-seq reveals a largely conserved genome organization relative to nuclear speckles with small position changes tightly correlated with gene expression changes. Genome Res. 31:251–264.

